# The phosphatase Glc7 controls eisosomal response to starvation via posttranslational modification of Pil1

**DOI:** 10.1101/2022.08.09.503340

**Authors:** Katherine M. Paine, Kamilla M. E. Laidlaw, Gareth J. O. Evans, Chris MacDonald

## Abstract

The yeast plasma membrane (PM) is organised into specific subdomains that regulate surface membrane proteins. Surface transporters actively uptake nutrients in particular regions of the PM where they are also susceptible to substrate induced endocytosis. However, transporters also diffuse into distinct subdomains termed eisosomes, where they are protected from endocytosis. Although most nutrient transporter populations are downregulated in the vacuole following glucose starvation, a small pool is retained in eisosomes to provide efficient recovery from starvation. We find the core eisosome subunit Pil1, a Bin, Amphiphysin and Rvs (BAR) domain protein required for eisosome biogenesis, is phosphorylated primarily by the kinase Pkh2. In response to acute glucose starvation, Pil1 is rapidly dephosphorylated. Enzyme localisation and activity screens implicate the phosphatase Glc7 is the primary enzyme responsible for Pil1 dephosphorylation. Both depletion of *GLC7* and phospho-ablative or phospho-mimetic mutations of Pil1 correlate with Pil1 phosphorylation status, failure to properly retain transporters in eisosomes, and results in defective starvation recovery. We propose precise posttranslational control of Pil1 modulates nutrient transporter retention within eisosomes depending on extracellular nutrient levels, to maximise recovery following starvation.

## INTRODUCTION

The plasma membrane (PM) of eukaryotic cells is organised into distinct domains of specific lipids and proteins (Kraft, 2013). In the budding yeast *Saccharomyces cerevisiae*, distinct spatiotemporal localisation patterns have been observed for different proteins (Berchtold and Walther, 2009; Grossmann et al., 2007; Heinisch et al., 2010; Murley et al., 2017; Spira et al., 2012). Original localisation studies distinguished between the hexose transporter Hxt1, that is uniformly dispersed across the surface and other proteins, such as Pma1 and Can1, that were found in discrete, non-overlapping regions (Malínská et al., 2003). The punctate subdomains occupied by the arginine transporter Can1, originally termed the Membrane Compartment of Can1 (MCC) and later denoted as eisosomes (Walther et al., 2006), have also been shown to house many other nutrient transporters (Babst, 2019). Eisosomes have been identified in other fungal species, such as *Aspergillus nidulans* and *Ashbya gossypii* (Seger et al., 2011; Vangelatos et al., 2010), as well as various species of lichens and algae (Lee et al., 2015).

Eisosomes are furrow-like PM structures enriched in sterols and sphingolipids (Grossmann et al., 2007; Malínská et al., 2003; StráDalová et al., 2009). Eisosome formation occurs *de novo*, and although once formed these structures are relatively immobile, core proteins remain dynamic (Moreira et al., 2009; Olivera-Couto et al., 2015; Walther et al., 2007). Many proteins localise to eisosomes, such as core structural proteins, post-translational modifiers, tetraspan membrane proteins and uncharacterised factors (Foderaro et al., 2017). For example, the tetraspanner Nce102, which functions as a sphingolipid sensor and promotes membrane curvature (Fröhlich et al., 2009; Haase et al., 2022; Vaskovicova et al., 2020; Zahumenský et al., 2022) and Seg1, a stability factor that operates upstream of eisosome formation (Moreira et al., 2012; Seger et al., 2011). A screen for PI(4,5)P_2_ regulators revealed eisosome factors Slm1 and Slm2 bind lipids, which are required for proper eisosomal organisation and integrate with TORC2 signalling and lipid synthesis (Audhya et al., 2004; Berchtold et al., 2012; Fadri et al., 2005; Kamble et al., 2011; Nagaya et al., 2002; Riggi et al., 2018). The Bin, Amphiphysin and Rvs (BAR) domain proteins Pil1 and Lsp1 are required for organising lipids during the sculpting of eisosomes (Moreira et al., 2009; Walther et al., 2006; Zhao et al., 2013; Ziólkowska et al., 2011). Pil1 and Lsp1 freely diffuse in the cytoplasm but almost exclusively localise to eisosomes at steady state (Olivera-Couto et al., 2015). Under stress conditions Lsp1 can at least partially complement loss of Pil1 (Vesela et al., 2023). Pil1 and Lsp1 are phosphorylated by kinases Slt2, Pkh1 and Pkh2, and the consequences of Pil1 phosphorylation on eisosome biogenesis have been characterized previously (Mascaraque et al., 2013; Walther et al., 2007; Zhang et al., 2004).

Yeast cells uptake nutrients from their external environment through specific transporters that localise to the PM (Jack et al., 2000; Léon and Teis, 2018). Regulation of these transporters at the PM allows for nutrient acquisition to be tightly controlled in response to cellular requirements. Active transporters localised to the PM, like Fur4, Can1 and Mup1, undergo conformational changes in response to nutrients and are more efficiently serviced by the endocytic machinery (Gournas et al., 2017; Guiney et al., 2016; Keener and Babst, 2013). Nutrient transporter ubiquitination is the signal for trafficking through the multivesicular body (MVB) pathway, where ubiquitinated proteins are recognised and packaged into intralumenal vesicles of the MVB by the endosomal sorting complex required for transport (ESCRT) apparatus (Migliano et al., 2022). Upon MVB-vacuole fusion, intraluminal vesicles containing surface proteins are deposited in the degradative environment of the vacuolar lumen. These feedback mechanisms allow transporter degradation to avoid excessive nutrient uptake, which can be detrimental (Kaur and Bachhawat, 2007; Séron et al., 1999; Watanabe et al., 2014). These trafficking events are coordinated in response to nutritional cues, for example in response to nitrogen starvation, surface proteins are degraded more readily by the elevation in vacuolar sorting triggered via Rsp5 and its adaptors (MacGurn et al., 2011; Müller et al., 2015). Rsp5 mediated degradation is also upregulated in response to growth past log-phase when niacin becomes limited (MacDonald et al., 2015). Surface proteins are also degraded faster and recycled less efficiently in response to leucine starvation (Jones et al., 2012; MacDonald and Piper, 2017). A similar dual control of trafficking pathways in response to glucose starvation, which triggers surface protein degradation (Lang et al., 2014), occurs through an increase in AP180 mediated endocytosis and a decrease in Gpa1-PI3-kinase mediated recycling (Laidlaw et al., 2021; Laidlaw et al., 2022b).

The tendency of nutrient transporters like Can1, Fur4 and Mup1 to also localise to eisosomes has led to various investigations into what regulatory control is provided within these subdomains (Busto et al., 2018; Gournas et al., 2018a; Gournas et al., 2018b; Moharir et al., 2018). Although activity of these transporters may vary when localised to eisosomes, the consensus view that eisosomes provide protection from ubiquitin mediated endocytosis has been demonstrated for all (Athanasopoulos et al., 2019). These surface cargoes have also been used in the context of stress condition experiments. Following stress, nutrient transporter populations are not entirely degraded, with a small proportion of the cellular pool being sequestrated in eisosomes. Unlike the response to substrate, where transporters like Can1, Fur4 and Mup1 move from eisosomes and undergo endocytosis, starvation conditions like poor nitrogen source or growth to stationary phase results in increased transporter concentration in eisosomes (Gournas et al., 2018b; Moharir et al., 2018). Stress conditions trigger restructuring of eisosomes, with changes in PM tension and deepening of these structures, to better retain this reserve pool of nutrient transporters (Appadurai et al., 2020; Moharir et al., 2018; Riggi et al., 2018). Furthermore, there is a physiological benefit to harbouring these nutrient transporters in eisosomes: to allow efficient recovery following a return to replete conditions. For example, the uracil transporter Fur4 that is required at the surface for efficient growth in limited uracil conditions contributes to efficient recovery following glucose starvation, in a uracil-dependent manner (Laidlaw et al., 2021; Paine et al., 2021).

The retention of nutrient transporters in response to stress is not well understood at a mechanistic level. As mentioned, phosphorylation of the core factor Pil1 by Pkh-kinases is an important regulatory step in eisosome biogenesis (Karotki et al., 2011; Luo et al., 2008; Walther et al., 2007). We find that under basal conditions, only Pkh2 predominantly localises to eisosomes and is largely responsible for phosphorylating Pil1.

As Pil1 is dephosphorylated in response to glucose starvation (Laidlaw et al., 2021), screening for responsible enzymes reveals the PP1 phosphatase Glc7 is a critical enzyme that regulates Pil1 dephosphorylation, nutrient transporter homeostasis and recovery from starvation. These effects are phenocopied upon mutation of Pil1 phospho-sites, suggesting that the phosphorylation status of Pil1 modulates eisosomes, not only during their biogenesis but to retain nutrient transporters at the surface for recovery following starvation.

## RESULTS

### Pkh2 is the predominant kinase that phosphorylates Pil1

It has been previously shown that Pil1 phosphorylation is ablated in a double *pkh1*^*ts*^ *pkh2Δ* mutant (Luo et al., 2008; Walther et al., 2007). To determine the contribution of Pkh1 and Pkh2 to the phosphorylation of Pil1, we assessed individual deletion mutants. As Pkh3 was identified as a multicopy suppressor *pkh*^*ts*^*pkh2* mutants (Inagaki et al., 1999) but has not been tested for a role in Pil1 phosphorylation, we included *pkh3Δ* mutants in this analysis. Pil1 phosphorylation status affects its migration during electrophoresis and can be visualised by immunoblot (Walther et al., 2007). Pil1 phosphorylation was assessed in all three *pkh* mutants to reveal *pkh2Δ* mutants were most defective and that *pkh1Δ* and *pkh3Δ* cells have only a small but significant defect (**Figures 1A and B**). We next performed localisation studies for these kinases. Although previous studies have demonstrated that over-expression of Pkh1 and Pkh2 with the galactose inducible promoter *GAL1* is required for sufficient levels to localise these proteins (Roelants et al., 2002; Walther et al., 2007), we avoided this glucose-repression strategy, due to its effect on eisosome biology (Laidlaw et al., 2021). GFP-tagged kinases were instead over-expressed from the constitutive *NOP1* promoter (Weill et al., 2018) in cells co-expressing the eisosomal marker Nce102 tagged with mCherry. Only GFP-Pkh2 predominantly colocalised with Nce102-mCherry (**Figures 1C and D**). This small apparent contribution of Pkh1 and Pkh3 to Pil1 phosphorylation observed by immunoblot might be explained by the fact that although most cells do not localise Pkh1 or Pkh3 to eisosomes, a small number of cells do (**Supplemental Figures S1A - S1B**). In further support of Pkh2 being a regulator of Pil1 phosphorylation, overexpressing Pkh2 leads to a significant increase in Pil1 phosphorylation observed in both wild-type and *pkh2Δ* cells (**Figures 1E and F**). These data demonstrate that Pkh2 is the primary Pkh-family member responsible for Pil1 phosphorylation, but that Pkh1 and Pkh3 also exhibit subsidiary roles.

**Figure 1:**
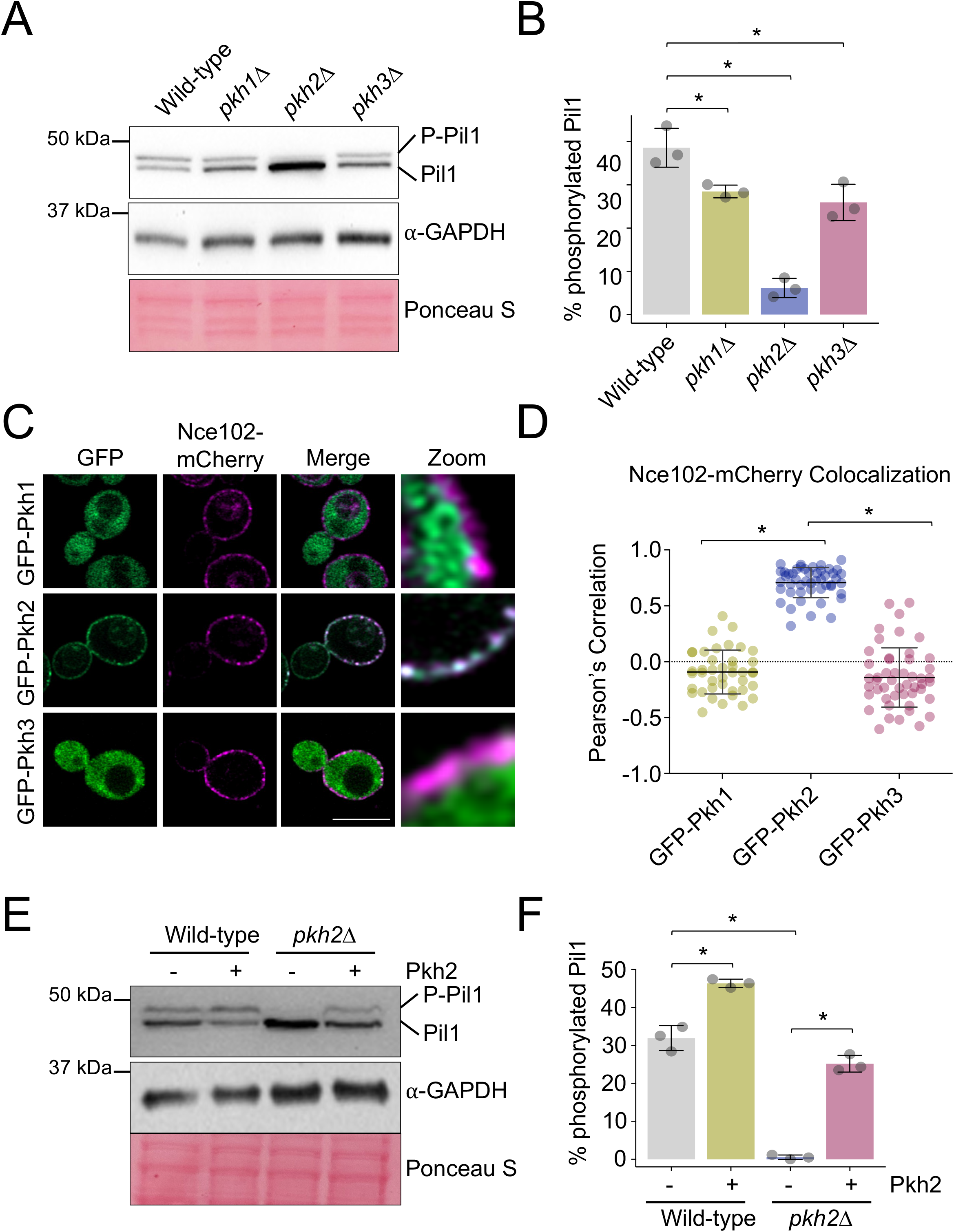
Pkh2 predominately regulates phosphorylation of Pil1. **A)** Whole cell lysates of wild-type, *pkh1Δ, pkh2Δ* and *pkh3Δ* cells were analysed by immunoblot using α-Pil1 and α-GAPDH antibodies, including Ponceau S stained membrane. **B)** The percentage phosphorylated Pil1 from each yeast strain was quantified (n=3). **C)** Cells co-expressing Nce102-mCherry and indicated GFP tagged Pkh-kinases were grown to mid-log phase and imaged using confocal microscopy (Airyscan 2). **D)** The Pearson’s Correlation Coefficient was measured between Nce102-mCherry with the respective GFP-tagged kinases (n > 40). **E)** Wild-type and *pkh2Δ* cells were transformed with either an empty vector control (-) or a 2μ Pkh2 over-expression plasmid (+). Whole cell lysates were generated from transformants and Pil1 phosphorylation assessed by immunoblot. Levels of GAPDH and Ponceau S are shown as loading controls. **F)** The percentage of phosphorylated Pil1 was quantified (n=3) and shown (right). Statistical significance indicated (*). Scale bar = 5μCm.

Phosphorylated peptides of Pil1 have previously been identified by mass spectrometry (Albuquerque et al., 2008; Luo et al., 2008; Swaney et al., 2013; Walther et al., 2007), suggesting multiple levels of potential phospho-regulation. Experimental work has determined several key phospho-sites (Luo et al., 2008; Walther et al., 2007), which map to distinct regions of Pil1 (**Figure 2A**). To ascertain if any additional kinases beyond the Pkh-family were responsible for Pil1 phosphorylation, NetPhorest analysis (Horn et al., 2014) was used to predict potential kinases for known Pil1 phospho-sites (**Figure 2B**). Mutants of any high scoring kinases (**Supplemental Table T1**) were tested for a role in phosphorylating Pil1 by immunoblot (**Supplemental Figure S2**). Only eight mutants showed any indication of a potential role, which was followed up quantitatively. This revealed *hog1*Δ cells lacking the yeast homologue of the mammalian MAPK p38, Hog1 (Han et al., 1994), had reduced levels of Pil1 phosphorylation. Also, depletion mutants with reduced levels of the Hippo-like kinase Cdc15 (Rock et al., 2013; Steensma et al., 1987), by virtue of a DAmP cassette (Breslow et al., 2008), showed higher levels of Pil1 phosphorylation (**Figure 2C**). This additional analysis again supports the notion that Pkh2 is the primary enzyme responsible for Pil1 phosphorylation, but that phosphorylation via other kinases may also have an impact, potentially through indirect mechanisms such as transcriptional or stress-induced control.

**Figure 2:**
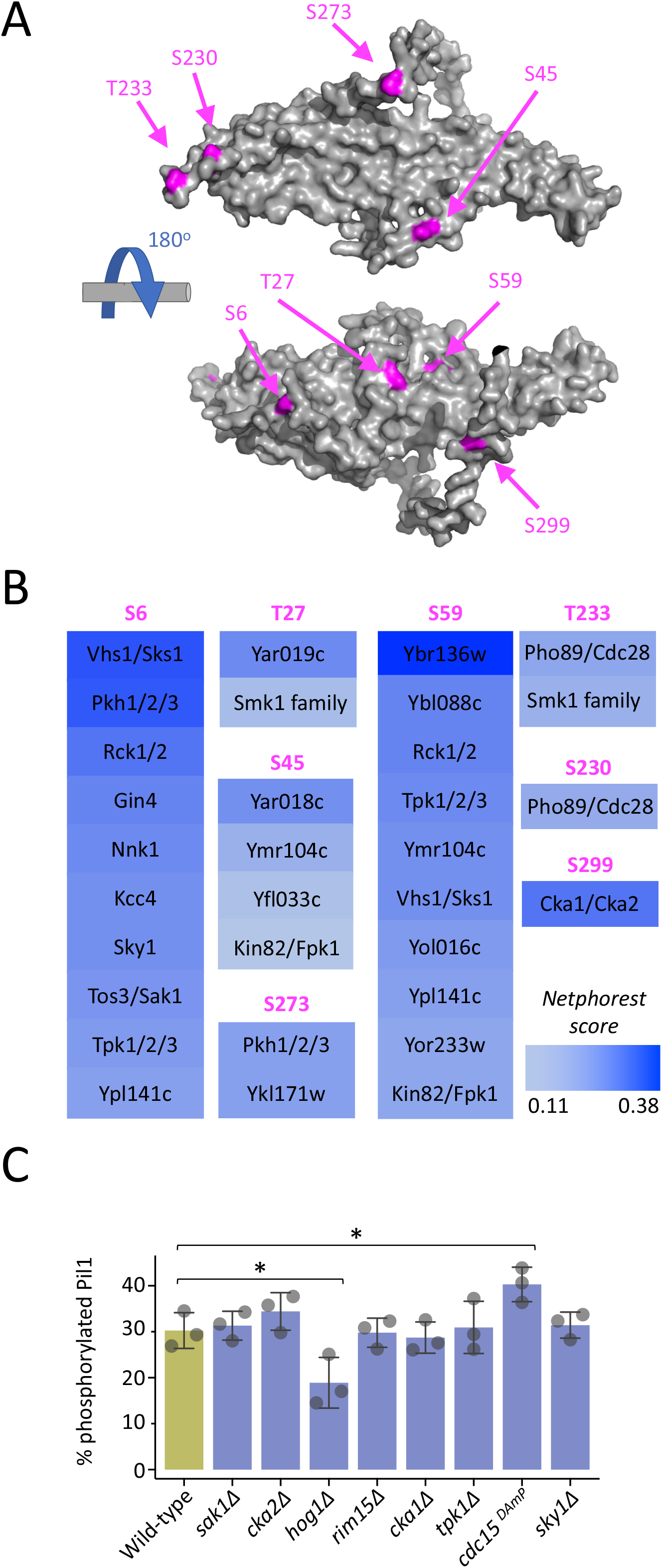
Bioinformatic screen for additional kinases that service Pil1. **A)** Alphafold structural model of Pil1 residues 1- 307 (grey) shown with eight previously verified phosphorylation sites indicated (magenta). **B)** The Pil1 protein sequence was surveyed using NetPhorest searching against a reference kinase database for *Saccharomyces cerevisiae*. Kinases that scored above threshold (0.1) are presented as a heat map (blue) with the indicated potential phosphorylated residue (magenta). **C)** Whole cell lysates of wild-type and kinase mutant cells were generated and percentage Pil1 phosphorylation assessed by immunoblot and presented as a histogram (n=3). Statistical significance indicated (*).

### Pil1 is dephopshorylated in response to glucose starvation

In response to acute glucose starvation, nutrient transporters localise to eisosomes (Laidlaw et al., 2021) and are hypothesised to relocate to PM regions for nutrient uptake upon return to replete conditions (**Figure 3A**). For glucose starvation, media lacking glucose but containing the trisaccharide raffinose, which cannot be quickly metabolised (de la Fuente and Sols, 1962), is used. Upon raffinose exchange, nutrient transporters such as Mup1 are primarily downregulated, however a small pool also concentrate within eisosomes (Laidlaw et al., 2021). This is best visualised by comparing puncta of Mup1-GFP in eisosomes marked by Pil1-mCherry in relation to Mup1 also diffusely localised to other PM regions from top focussed confocal slices (**Figures 3B**). During the initial period of glucose starvation when nutrient transporters accumulate in eisosomes, we observe rapid Pil1 dephosphorylation (**Figure 3C - 3D**). We also observe significant glucose induced dephosphorylation in *pkh1Δ*, moving to levels similar to *pkh2Δ* in basal conditions, again suggesting Pkh1 is not a key regulator of Pil1 (**Supplemental Figures S3A - S3B**).

**Figure 3:**
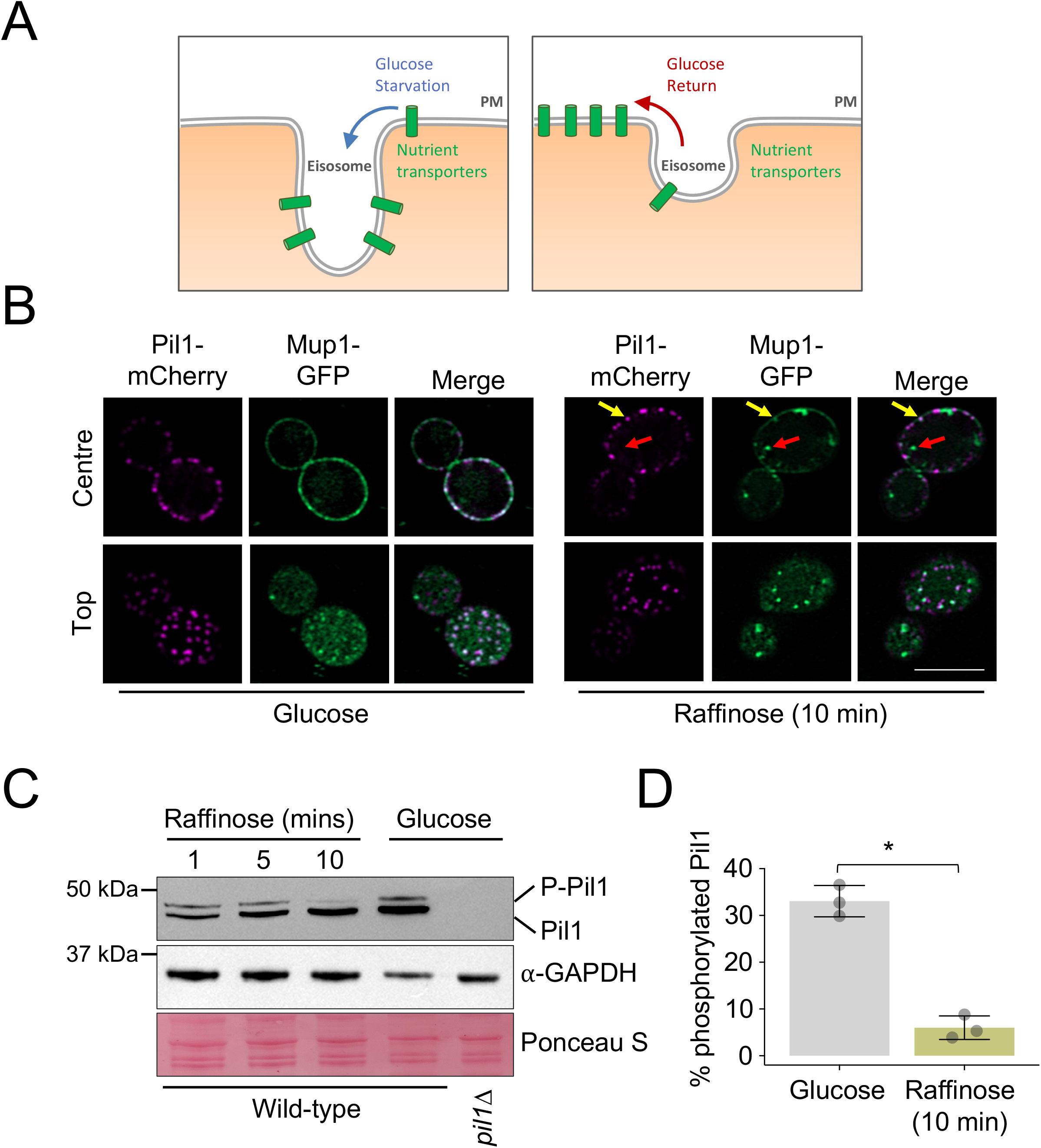
Pil1 is dephosphorylated in response to glucose starvation. **A)** Schematic showing the increased diffusion of nutrient transporters into eisosomes in response to glucose starvation, and their potential exit to aid recovery in replete conditions. **B)** Cells co-expressing Pil1-mCherry and Mup1-GFP were imaged using confocal microscopy (Airyscan 2) with centre and top focus under glucose conditions and following 10 minutes of exchange with raffinose media. Mup1 localised to endosomes (red arrow) and Pil1 marked eisosomes (yellow arrow) after raffinose treatment are indicated. **C)** Wild-type cells exposed to raffinose media for 1, 5 and 10 minutes prior to lysate generation were immunoblotted using α-Pil1 antibodies and compared to wild-type and *pil1Δ* cells grown in glucose replete conditions. GAPDH blot and Ponceau S stained membrane is included as a loading control. **D)** Percentage of phosphorylated Pil1 was generated for WT vs 10 minutes of raffinose treatment for wild-type cells was quantified (right). Statistical significance indicated (*). Scale bar = 5μCm.

As transporters are sequestered in eisosmes whilst Pil1 is dephosphorylated in response to glucose starvation, we hypothesise that Pil1 dephosphorylation plays a role in retaining nutrient transporters. To test this idea, we first set out to identify any responsible phosphatase enzymes and then test if they have an impact on eisosomes, nutrient transporters, or starvation recovery. In *Saccharomyces cerevisiae* 43 phosphatases have been identified (Offley and Schmidt, 2019), of which 39 are non-essential and 4 are essential. To identify the phosphatase(s) responsible for Pil1 dephosphorylation, we screened mutants of all phosphatase enzymes for their activity in glucose and raffinose conditions (**Figure 4A**). Pil1 phosphorylation status was assessed in null mutants (*Δ*) lacking non-essential phosphatases or with reduced expression of essential phosphatases, by virtue of a DaMP cassette (Breslow et al., 2008). Mutants were scored based on defects in Pil1 dephosphorylation (**Figure 4B**). This screen revealed 7 top scoring phosphatase mutants selected for further quantitative analysis (**Figure 5A - 5B**). Next, we assessed the localisation of many GFP tagged phosphatases at mid-log phase and stationary phase (**Figure 5C, Supplemental Figure S4**). We included stationary phase as a nutritional stress associated with eisosome transporter retention (Gournas et al., 2018b). This confirmed a range of localisations, many to the nucleus and cytoplasm but also Ppn2 at the vacuole and Ptc5 at the mitochondria (Breker et al., 2013). Interestingly, we also observed several GFP tagged phosphatases that had changes in localisation following growth to stationary phase, including Sdp1, Ptc5, and Nem1. Although GFP-Msg5 and GFP-Siw14 fusions localised to the periphery upon growth to stationary phase, deletion of these mutants had no impact on Pil1 phosphorylation, so we assume this peripheral localisation is not related to eisosomal regulation. However, GFP-Glc7 showed significant peripheral punctate localisation in both growth conditions, in addition to localisation to the mid-body, the cytoplasm and the nucleus (Bloecher and Tatchell, 2000; Breker et al., 2013). The only phosphatase that showed significant localisation to the cell periphery and that exhibited defects in phosphorylation upon mutation was Glc7.

**Figure 4:**
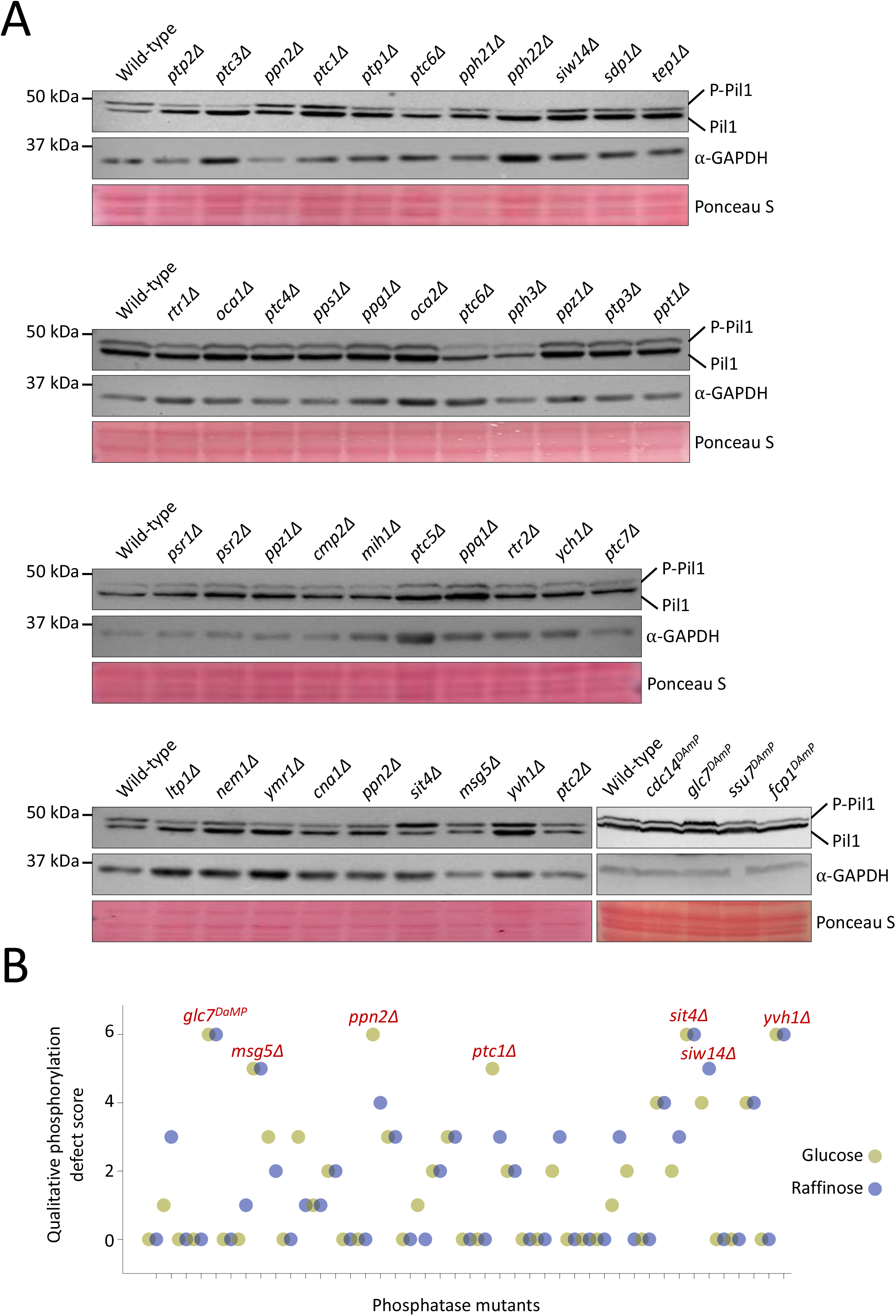
Primary activity screen for regulators of Pil1 dephosphorylation. **A)** Wild-type cells and indicated phosphatase mutants were grown to log phase in glucose replete conditions prior to lysate generation and immunoblotting with α-Pil1 and α-GAPDH antibodies. Representative blots for at least one mutant are shown, alongside a Ponceau S stained membrane. **B)** Immunoblots for all phosphatase mutants were qualitatively scored based on their Pil1 phosphorylation phenotype in both glucose (green) and raffinose (purple) conditions compared with wild-type controls. The highest scoring mutants are indicated in red text.

**Figure 5:**
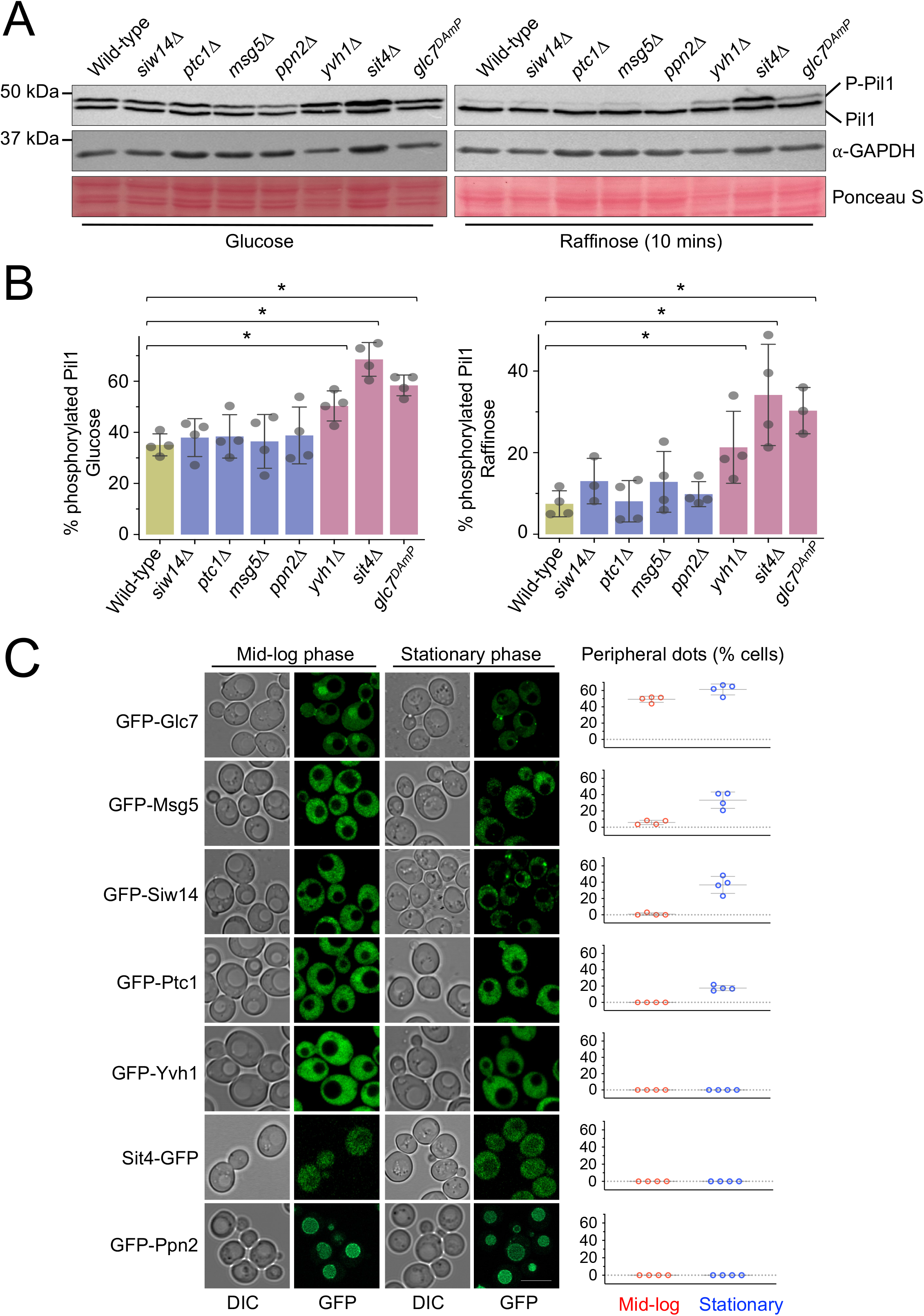
Secondary activity and localisation screens of phosphatase candidates. **A)** Whole cell lysates of wild-type and phosphatase mutant candidates from glucose replete media (left) or following 10 minutes exchange with raffinose media were generated and analysed by immunoblotting using α-Pil1and α-GAPDH antibodies. Ponceau S stain shown as an additional loading control. **B)** Quantification of percentage Pil1 phosphorylated in indicated mutants was calculated. **C)** Yeast expressing GFP tagged phosphatases were cultured to mid-log and stationary phase prior to confocal microscopy (Airyscan). The number of peripheral dots per cell (n > 30) from separate experiments (n = 4) were quantified in each condition (right). Statistical significance indicated (*). Scale bar = 5 μCm.

### Glc7 controls dephosphorylation of Pil1

Glc7 is an essential phosphatase (Clotet et al., 1991; Feng et al., 1991) that has been previously shown to function in glucose related pathways, where it acts with its regulatory subunit Reg1 (Tu and Carlson, 1995) and bud neck formation (Larson et al., 2008), amongst other roles in the cell. We confirmed that *glc7*^*DAmP*^ results in reduced dephosphorylation of Pil1 in both glucose replete and glucose starvation conditions (**Figures 6A and B**). As a complementary approach to test the role of Glc7 in Pil1 phosphorylation we altered *GLC7* expression levels using a YETI (Yeast Estradiol with Titratable Induction) strain (Arita et al., 2021), which allows modulation of expression by varying ß-estradiol concentrations (**Figure 6C**). We first show that protein levels of Glc7 can be controlled by exogenous ß-estradiol concentrations (**Figure 6D**), and this is specific to the *YETI-GLC7* strain (**Supplemental Figures 5A - 5B**). We then demonstrate that altering Glc7 levels, using a gradient of 0 nM, 12.5 nM and 100 nM ß-estradiol, correlates with Glc7 activity in dephosphorylating Pil1 in a concentration dependent manner (**Figures 6E - 6F and Supplemental Figure S5C**). We then tested if GFP-Glc7 localised to eisosomes marked with Pil1-mCherry. Although, neither steady state or time lapse imaging in glucose and raffinose media revealed large amounts of Glc7 localisation to eisosomes, there were often small regions of colocalisation at some eisosomes (**Figure 6G and Supplemental Figure S5D**). This combinatorial approach strongly suggests that Glc7 is responsible for dephosphorylation of Pil1, and this might be achieved via Glc7 directly at eisosomes.

**Figure 6:**
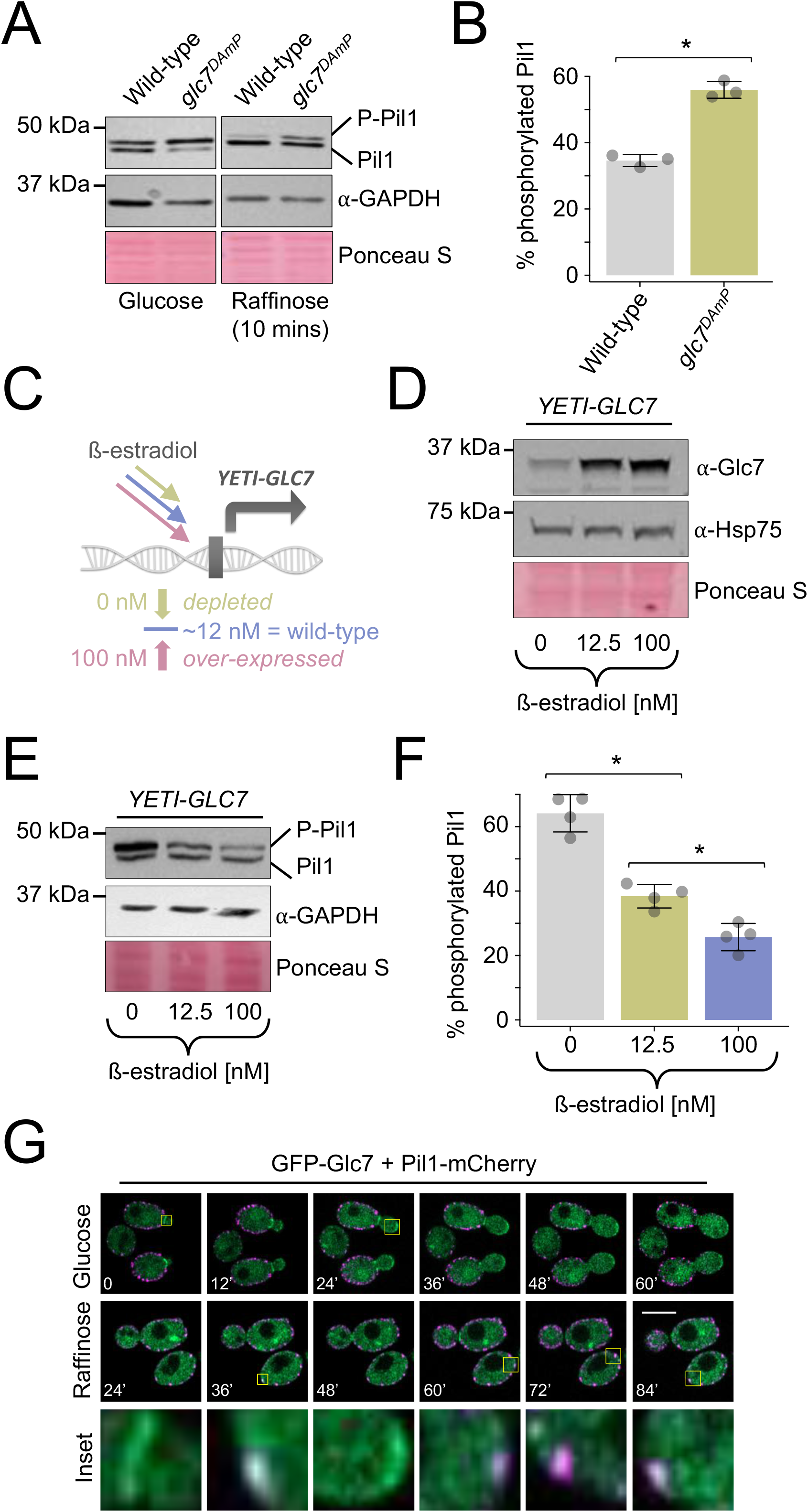
Glc7 mediates dephosphorylation of Pil1. **A)** Wild-type and *glc7*^*DAmP*^ cells were cultured in glucose media and following 10 minutes of raffinose treatment prior to the generation of whole cell lysates and immunoblotting with α-Pil1 and α-GAPDH antibodies. Ponceau S stain shown as an additional loading control. **B)** The percentage of phosphorylated Pil1 from A was quantified (n=3). **C)** Schematic outlining the principle of the YETI expression system for titratable expression of *GLC7* to mimic severely depleted (green) and over-expressed (pink) levels. **D - E)** *YETI-GLC7* cells were grown overnight in 12.5 nM β-estradiol before washing ×3 in YPD media and dilution in fresh media containing 0, 12.5 and 100 nM β-estradiol. Cells were then grown for 6 hours prior to the generation of whole-cell lysates and immunoblots using **D)** α-Glc7 and **E)** α-Pil1 antibodies. Loading controls using α-GAPDH antibodies and Ponceau S stained membrane are included for each. **F)** Pil1 phosphorylation was quantified (n=3) from β-estradiol titrations shown in (**E**). **G)** Time-lapse microscopy of GFP-Glc7 and Pil1-mCherry expressing cells was performed in glucose and following 25 minutes raffinose, with indicated time slices labelled. Statistical significance indicated (*). Scale bar = 5μCm.

### Phosphorylation of Pil1 is important for starvation recovery

Having implicated Glc7 in Pil1 dephosphorylation, which occurs following glucose starvation, we next wanted to test if this was connected to nutrient transporter residence in eisosomes and starvation recovery. Previous studies have assessed the phosphorylation profile of various Pil1 mutants with mutation of 8 verified phospho-sites (Luo et al., 2008; Walther et al., 2007). We generated phospho-ablative (changed to alanine, 8A) and phospho-mimetic (changed to aspartate, 8D) versions of Pil1 at these phospho-sites (**Figure 7A**). Western blotting confirmed that the 8A and 8D mutations resulted in Pil1 migrating not as a doublet, but as a single band, with faster migration of the phospho-ablative Pil1-8A-mGFP and slower migration of the phospho-mimetic Pil1-8D-mGFP fusion (**Figure 7B**). Fluorescence microscopy of these mGFP tagged Pil1 versions showed both 8A and 8D expressing strains exhibited an altered localisation phenotype compared to wild-type cells (**Figure 7C**), as previously documented for phospho-mutants of Pil1 (Luo et al., 2008; Walther et al., 2007). We quantified these differences (**Supplemental Figure S6**), revealing both mutants have fewer eisosomes compared to wild type (**Figure 7D**) in addition to more cytoplasmic signal (**Figure 7E**). This analysis showed a more pronounced defect in eisosome number and levels for Pil1-8D-mGFP than Pil1-8A-mGFP. To understand how nutrient transporter localisation might be affected in strains expressing phospho-mutants, we expressed the arginine transporter Can1, a dual reporter for eisosome morphology and transporter localisation. As expected, the abnormal localisation of Pil1-mGFP phospho-mutants is mirrored by Can1-mCherry, with fewer eisosome foci that colocalise with Pil1-8A and Pil1-8D (**Figure 7F**). Although Can1-mCherry still localises to the PM when Pil1-8A and Pil1-8D are expressed, the punctate eisosome pattern is reduced and there is a small amount mis-localised to the vacuole (discussed in next section).

**Figure 7:**
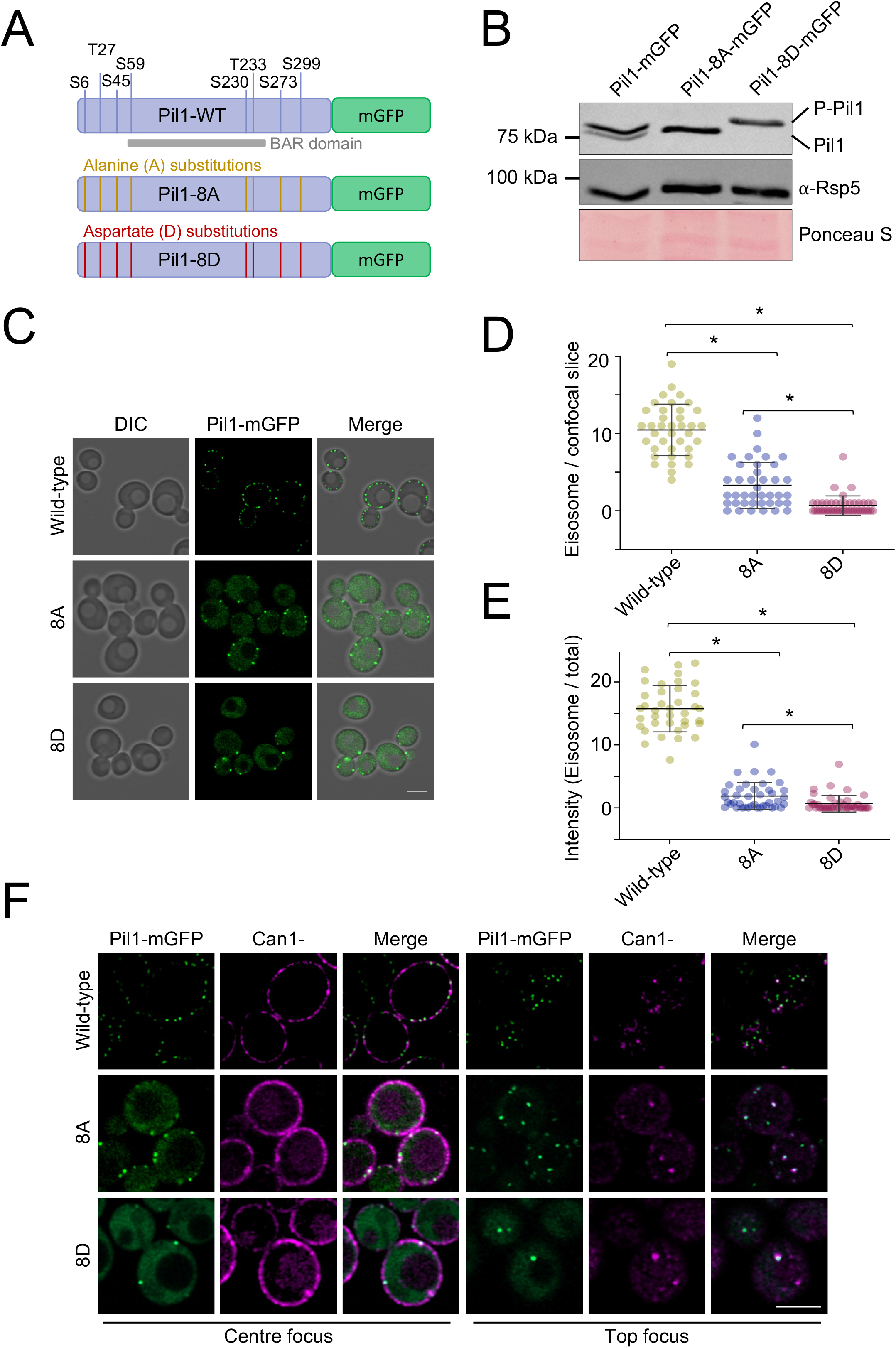
Characterisation of Pil1 phospho-mutants. **A)** Illustration showing Pil1 fusion to mGFP, including its verified phospho-sites and BAR domain. Pil1 cassettes that have been mutated to alanine (yellow) or aspartate (red) that were stably integrated at the *PIL1* locus are also shown. **B)** Pil1 or indicated 8A and 8D mutants were expressed from the endogenous locus as mGFP fusions in strains grown to mid-log phase, harvested and lysate generated for immunoblotting with α-Pil1 and α-GAPDH antibodies. Ponceau S stain included as an additional control. **C)** Versions of mGFP tagged Pil1 were expressed as the sole chromosomal copy and localised by confocal microscopy coupled to Airyscan 2 detector. **D)** Pil1 labelled eisosomes were identified by otsu segmentation and number per centre focussed confocal slice quantified (n = > 37). **E)** Integrated density for all GFP tagged Pil1 versions localised to eisosomes identified from segmentation performed in (**D**) was calculated as a percentage of the total signal. **F)** Wild-type and the Phospho-mutants expressing Can1-mCherry were imaged using confocal microscopy coupled to Airyscan2. Micrographs show centre and top focus. Statistical significance indicated (*). Scale bar = 5μCm.

To test if Pil1 phospho-mutants were functional in starvation recovery, we used an assay that monitors recovery growth (Laidlaw et al., 2021). Although a subtle growth defect is observed for both Pil1-8A and Pil1-8D in rich media, growth was indistinguishable to wild-type cells in SC minimal media (**Supplemental Figure S7A**). All cells were cultured to log phase in SC media followed by 2-hours glucose starvation. Upon a return to replete SC media, we find that both Pil1-8A and Pil1-8D mutations are defective in recovery growth compared with wild-type cells (**Figure 8A - 8B**). To test if these results can be explained in the context of deregulation of nutrient transporters via Glc7 activity, we expressed Can1-mCherry in wild-type and *glc7*^*DAmP*^ cells. These imaging results showed eisosome biogenesis still occurs in cells with less Glc7, but there was a significant reduction in eisosme number (**Figure 8C - 8D**). Increased contrast of micrographs and quantification revealed that most *glc7*^*DAmP*^ cells expressing Can1-mCherry, mis-localise a portion to the vacuole (**Figures 8C and 8E**), much like phospho-mutant versions of Pil1. We also assessed Can1-mCherry mis-localisation biochemically, by immunoblotting cells and measuring stable mCherry following proteolytical processing in the vacuole. In agreement with the micrographs, essentially no Can1-mCherry is found in the vacuole in wild-type cells but a significant intravacuolar population is observed upon depletion of Glc7 (**Figures 8F - G**). In support of our hypothesis that the described effects on eisosomes and nutrient transporter trafficking caused by depletion of Glc7 affect physiology following starvation, we used the recovery assay to show that despite no growth defects of *glc7*^*DAmP*^ (**Supplemental Figure S7B**), the ability of *glc7* mutants to recover efficiently from glucose starvation is hampered (**Figures 8H - I**).

**Figure 8:**
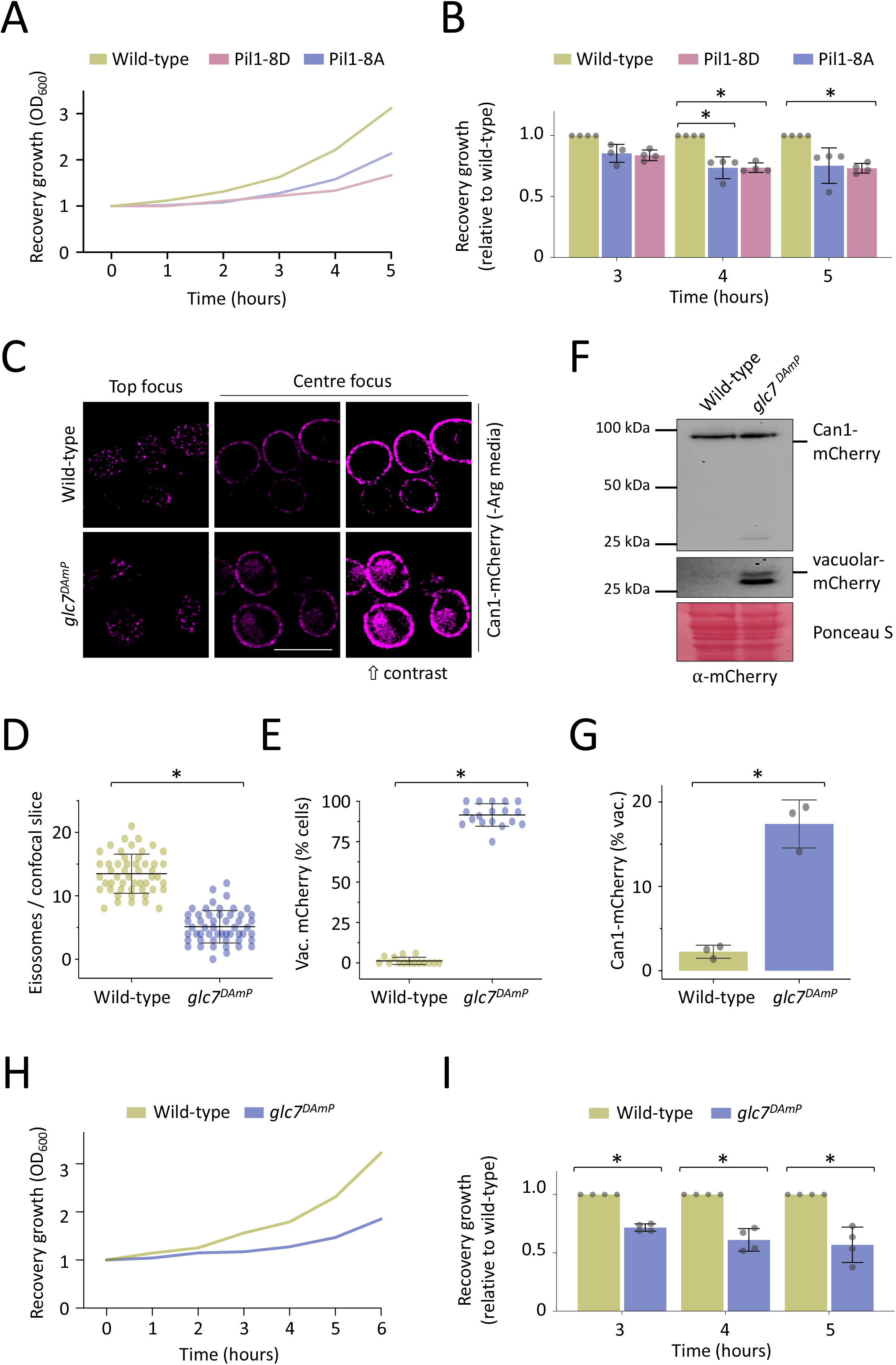
Glc7 is required for efficient recovery from glucose starvation. **A)** Wild-type and phospho-mutant strains were grown to log phase in SC minimal media, equivalent cell numbers were estimated by OD_600_ measurements and harvested. 10-fold serial dilutions were generated, and yeast spotted out on SC plates. Growth was recorded at 24 and 48 hours. **B)** Cells at mid-log phase were subjected to 2 hours of glucose starvation (raffinose treatment), returned to glucose replete conditions and growth measured over time. **C)** The growth assay in (**B**) was repeated (n = 4) and the growth relative to wild-type was quantified for each indicated time-point. **D)** Wild-type and *glc7*^*DAmP*^ cells expressing endogenously expressed Can1-Cherry were grown in SC media lacking arginine and imaged using confocal microscopy (Airyscan 2). Micrographs show top and centre focus, with increased contrast of the latter to show intravacuolar signal. **E)** Eisosomes from top focussed confocal slices were quantified. **F)** Cells exhibiting intravacuolar mCherry signal were quantified from centre focussed micrographs. **G)** Wild-type and *glc7*^*DAmP*^ cells expressing Can1-mCherry were grown to mid-log phase, harvested and lysates were generated before immunoblotting using α-mCherry antibodies. Full length Can1-mCherry and vacuolar processed mCherry fragment are indicated, including enhanced exposure of the latter. Ponceau S stain included as a loading control. **H)** Densitometry was used to quantify the amount of vacuolar processed mCherry from experiemnts in (**G**). **I)** Wild-type and *glc7*^*DAmP*^ strains were grown to log phase, equivalent cell numbers were estimated by optical density measurements and harvested. 10-fold serial dilutions were generated, and yeast spotted on SC plates followed by recording growth at 24 and 48 hours. **J)** Cells at mid-log phase were subjected to 2 hours of glucose starvation (raffinose treatment) and then returned to glucose replete conditions, returned to glucose replete conditions and growth measured over time. **K)** The growth assay in (**J**) was repeated (n = 4) and growth relative to wild-type was quantified for indicated time-points. Statistical significance indicated (*). Scale bar = 5μCm.

Catalytic subunits of Type 1 protein phosphatase (PP1) enzymes are known to co-function with a plethora of non-catalytic subunits, which can regulate specificity and localisation of enzymes (Virshup and Shenolikar, 2009). The best characterised regulatory subunit of Glc7 is Reg1, which is required for glucose repression (Tu and Carlson, 1995). Reg1 also has a paralogue Reg2, that is not involved in glucose repression but functionally complements the growth defect of *reg1Δ* cells (Frederick and Tatchell, 1996). However, deletion of either *REG1* or *REG2* alone or in combination had no significant effects on Pil1 phosphorylation (**Figures 9A - B**). Genetic approaches have identified many factors thought to co-function with Glc7, with several predicted to be regulatory subunits (Logan et al., 2008). However, genetic mutation of these nine factors did not affect Pil1 phosphorylation (**Supplemental Figure S8A - S8B**). We then speculated that if an unknown factor existed, it might physically bind both Pil1 and Glc7. Bioinformatics revealed various candidates including some known to localise to the periphery (**Figure 9C**). However, although Ygr237c localises quite prominently to eisosomes (**Supplemental Figure S8C**), deletion of these factors also failed to identify a Pil1 regulator in either glucose or raffinose conditions (**Figures 9D - 9G)**.

**Figure 9:**
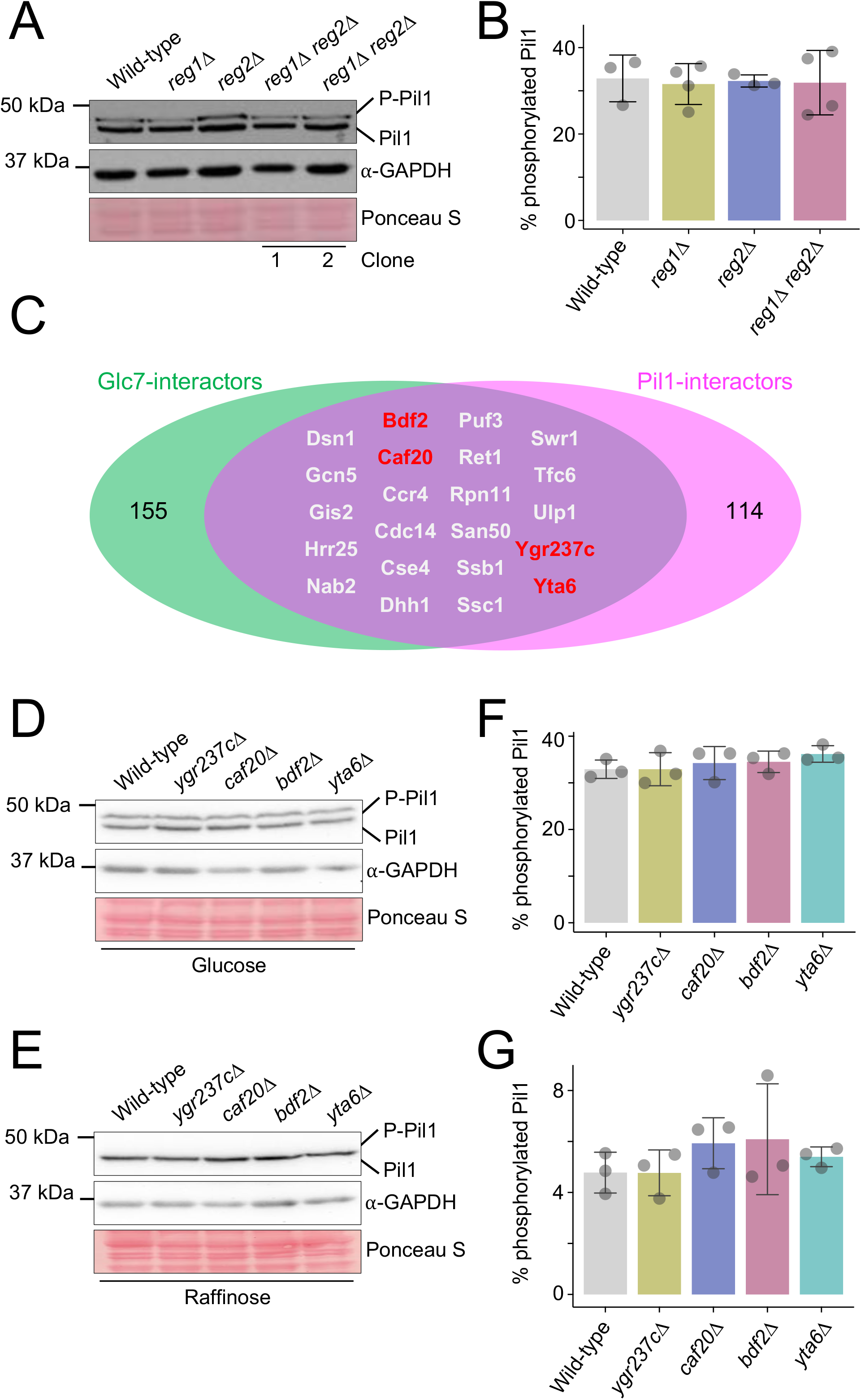
Glc7 regulatory subunit Reg1 does not appear to be involved in this response. **A)** Wild-type cells and indicated mutants were grown to log phase in glucose replete conditions prior to lysate generation and immunoblotting with α-Pil1 and α-GAPDH antibodies. Ponceau S stained membrane included. **B)** The percentage phosphorylated Pil1 from each yeast strain was quantified (n ≥ 3). **C)** Venn diagram showing the overlap of Glc7 and Pil1 interactors. **D - E)** Wild-type cells and indicated mutants were grown to log phase in glucose replete (**D**) or subjected to 10 minutes raffinose treatment (**E**) conditions before equivalent volumes were harvested for immunoblotting with α-Pil1 and α-GAPDH antibodies. **F - G)**. The percentage phosphorylated Pil1 from each yeast strain (**D - E**) was quantified (n = 3). Statistical significance indicated (*).

## DISCUSSION

Organisation of the yeast PM is complex with various surface localisation patterns known for both integral membrane proteins and surface associated factors (Spira et al., 2012). Since the discovery of eisosome subdomains, progress has been made in understanding the formation and biological function of these structures, particularly in response to cellular stress (Babst, 2019; Megarioti et al., 2023; Moseley, 2018). The discovery that Pil1 and Lsp1, and their phosphorylation by the Pkh-family kinases, are required for proper eisosome biogenesis (Walther et al., 2007; Walther et al., 2006) suggests post-translational modification of core components could regulate the eisosome environment. Pkh1 and Pkh2 were originally identified as homologues of human and *Drosophila* 3-phosphoinositide-dependent protein kinase-1 (PDK1) that are essential for viability (Casamayor et al., 1999). As this study demonstrated, the double *pkh1Δ pkh2Δ* yeast mutant was inviable, a strain harbouring a temperature sensitive allele of *PKH1* (D398G) and deletion of *PKH2*, (termed *pkh1*^*ts*^ *pkh2Δ*), was used to study kinase signalling pathways (Inagaki et al., 1999). This double *pkh1*^*ts*^ *pkh2Δ* mutant also revealed an early association of Pkh-kinases with endocytosis, as internalization from the PM was impaired (Friant et al., 2001). Due to the shared essential function of Pkh1 and Pkh2, the double mutant was the most logical strain to test effects on biogenesis of eisosomes (Walther et al., 2007). However, we clarify that Pkh2 is predominantly responsible for phosphorylating Pil1 with only minor roles for Pkh1 and the related kinase Pkh3, which also do not significantly localise to eisosomes (**Figure 1**). One key difference to our work and a previous study that found Pkh1 at eisosomes (Walther et al., 2007) is that we did not use the *GAL1* promoter for over-expression, so it may be that the glucose starvation stress of galactose induction media alters Pkh1 localisation or the eisosome environment. Beyond this, Pkh2 expressed from its endogenous promoter is also found at eisosomes (Fröhlich et al., 2009). Exploring other potential kinases and phospho-sites did reveal the Hog1 and Cdc15 kinases might also exhibit small roles regulating Pil1, but these effects were also relatively modest compared with Pkh2.

The finding that extracellular stress results in the accumulation of nutrient transporters in eisosome compartments, which deepen to facilitate this process, suggests a key role of eisosomes is related to nutrient uptake following stress (Appadurai et al., 2020; Gournas et al., 2018b). We have previously shown acute glucose starvation (2 hours) results in concentration of the nutrient transporter Mup1 to eisosomes. As eisosomal mutants fail to properly retain nutrient transporters and fail to recover efficiently from starvation (Laidlaw et al., 2021), we propose acute glucose starvation modulates eisosomes to better harbour transporters for recovery. In this study, we show the core eisosomal Pil1 is rapidly dephosphorylated during this acute glucose starvation period, and systematic screening of enzyme activity and localisation identified the PP1 phosphatase Glc7 as a Pil1 modifier (**Figure 3 - 5**). This led to the model that the phosphorylation status of Pil1 is important for reorganisation of existing eisosomes during cargo retention, in addition to its established role in eisosome biogenesis. As membrane bending effects of other BAR domain proteins are known to be affected by disordered regions (Busch et al., 2015; Zeno et al., 2018), it is conceivable that the charge / phosphorylation status of Pil1 modulates its lipid binding / sculping capacity, leading to nutrient transporter retention following starvation.

We also note that our screen identified significant effects on Pil1 dephosphorylation in both *sit4Δ* and *yvh1Δ* null strains. Sit4 is a PP2A type phosphatase first identified as regulating the cell cycle (Ronne et al., 1991). Since then it has many functional associations with nutrient signalling (Conrad et al., 2014). Sit4 has also been implicated in trafficking pathways used by nutrient transporters, at the endoplasmic reticulum (Bhandari et al., 2013) and from endosomes to the surface (Amoiradaki et al., 2021) and vacuole (Gurunathan et al., 2002). Yvh1 is a phospho-tyrosine specific enzyme (Guan et al., 1992) that is mainly associated with ribosome maturation (Kemmler et al., 2009) but has been implicated in autophagy downstream of TORC1 following nutrient depletion (Yeasmin et al., 2015). Although beyond the scope of this work, it will be interesting to test if these enzymes contribute to Pil1 dephosphorylation in response to nutrient starvation, either directly or indirectly. However, our screens identified the essential Type 1 Serine/Threonine protein phosphatase Glc7 (Cannon et al., 1994; Peng et al., 1990) as the most likely candidate to regulate eisosomes via Pil1. Glc7 has several roles in the cell including, growth, mitosis, transcription, stabilisation of emerging buds, glycogen metabolism and ion homeostasis (Feng et al., 1991; Hisamoto et al., 1995; Kozubowski et al., 2003; Peggie et al., 2002; Sanz et al., 2004; Williams-Hart et al., 2002), which we corroborate with expected localisations at the bud-neck, nucleus and cytoplasm (**Figure 5C**). Our imaging experiments showed some occasions where Glc7 localises to Pil1 at eisosomes (**Figure 6G**), so it may be that at steady state only a small percentage of Glc7 is required for eisosome maintenance, or this association may be transient. We confirmed Glc7 regulates Pil1 by depleting Glc7 using a Decreased Abundance by mRNA Perturbation (*DAmP*) method (Breslow et al., 2008) in addition to a Yeast Estradiol with Titratable Induction (YETI) depletion strategy (Arita et al., 2021), both of which showed elevated levels of Pil1 phosphorylation (**Figure 6**). This latter approach allowed fine tuning of Glc7 protein levels, which coincided with predicted Glc7 activity levels. We have recently also used the YETI system to produce null, wild-type and mutant levels of the endosomal protein Ist1 (Laidlaw et al., 2022a). As YETI depletion / over-expression effects are induced within 6 hours, this is an exciting alternative to genetic perturbations like CRISPR / recombination induced nulls, which take many generations to isolate a clonal population, allowing ample time for compensation to occur (El-Brolosy and Stainier, 2017).

As our phospho-ablative and phospho-mimetic versions of Pil1 were both defective, we assume being locked in either biochemical state does not promote eisosomal retention of transporters, but rather the fine-tuning of Pil1 phosphorylation is required to better harbour transporters acutely in response to nutritional stress. This mechanism could be important for understanding metabolic response of yeast to varying nutrient conditions, including pathogenic fungi (Rutherford et al., 2019). Glc7 having a role in eisosomal modulation during glucose starvation is conceptually consistent with many studies demonstrating that Glc7 integrates with transcriptional repression in response to glucose availability, via Snf1 and downstream factors (Sanz et al., 2000; Tu and Carlson, 1994). However, as the changes we observe in both Pil1 dephosphorylation and transporter retention are very rapid, we assume these effects are not mediated at the transcriptional level. Other eisosome based regulation is known to span much longer periods of starvation (Gournas et al., 2018b; Megarioti et al., 2023) and likely to have transcriptionally based control. In further support of this being a novel and distinct role for Glc7, the regulatory subunit Reg1, which is required for many transcriptional related Glc7-activities (Alms et al., 1999; Cui et al., 2004; Dombek et al., 1999), or its paralogue Reg2 (Frederick and Tatchell, 1996), showed no increase in phosphorylated Pil1 species upon deletion (**Figure 9**). Our additional efforts, both based on targets identified from the literature and through our own bioinformatic approaches, did not identify any other regulatory subunits of Glc7 involved with modifying Pil1. This might be explained by the fact that Glc7 has 100s of potential regulators that have yet to be characterised (Logan et al., 2008; Ramaswamy et al., 1998), and the hypothetical possibility that some are functionally redundant.

Nonetheless, the fact that Glc7 is robustly associated with glucose metabolism via distinct mechanisms suggests there may be more complexity to cellular response following starvation. It also remains to be understood how Glc7 senses glucose starvation prior to modifying eisosomes. One intriguing hypothesis is Glc7 activity is mediated via Pmp3, a cell periphery protein that is involved in maintenance of PM potential (Navarre and Goffeau, 2000). *pmp3Δ* mutants have defects in Pil1 phosphorylation and altered nutrient transporter stability, leading to the idea that Pmp3 is a phosphoinositide-regulated stress sensor (De Block et al., 2015). If so, this function may integrate with Glc7 modification of Pil1 following starvation, an idea supported by a genetic interaction between Glc7 and Pmp3 (Costanzo et al., 2010). Such mechanisms for eisosome regulation might also functionally connect to lipid homeostasis mediated by TORC2 (Riggi et al., 2018). This described role of stimuli induced post-translational modification of lipid binding proteins, which triggers remodelling of membranes, might also apply to other compartments and other eukaryotic systems.

## METHODS

### Reagents

**Supplemental Table T2** documents yeast strains used in this study.

### Cell culture

Yeast cells were routinely grown in YPD (1% yeast extract, 2% peptone, 2% dextrose) or synthetic complete (SC) minimal media (2% glucose, 0.675% yeast nitrogen base without amino acids, plus appropriate amino acid dropouts for plasmid selection) (Formedium, Norfolk, UK). 2% Glucose was routinely used, where stated 4% glucose was used. Cells were subjected to glucose starvation using 2% raffinose rather than glucose as described previously (Laidlaw et al., 2021). Plasmid pCM1054 is a 2μ over-expression plasmid for Pkh2 (Jones et al., 2008) used in **Figure 1E**.

### Mating of yeast strains

Single colony haploid BY4741 mat **a** yeast strains encoding *URA3-*GFP-tagged genes (Weill et al., 2018) and BY4742 Mat **α** modified at indicated loci with mCherry-his5^+^ cassettes were isolated on YPD agar. Strain isolates were then mixed and mated on YPD agar, then single diploid colonies were isolated on SC media lacking uracil and histidine, prior to confirmation of co-expression by fluorescence microscopy.

### Immunoblotting

Strains were grown to mid-log phase and equivalent volumes were harvested or starved for glucose with raffinose treatment prior to harvesting. Cells were treated to 0.2 N NaOH for 5 minutes prior to resuspension in lysis buffer (8 M urea, 10% glycerol, 50 mM Tris-HCl pH 6.8, 5% SDS, 0.1 % bromophenol blue and 10% 2-mercaptoethanol). SDS-PAGE was used to resolve proteins which were then transferred to a nitrocellulose membrane using the iBlot dry transfer system (Invitrogen). Ponceau S strain was used to confirm successful transfer and equal loading. Membranes were probed with antibodies stated, details listed in (**Supplemental Table T3**), and visualised using enhanced chemiluminescence (ECL) Super Signal Pico Plus (Thermo) and captured using a ChemiDoc Imager (Bio-Rad).

### Confocal microscopy

Yeast cells expressing fluorescently tagged proteins were grown to mid-log phase (unless stated) and then visualised in minimal media at room temperature on Zeiss laser scanning confocal instruments (Zeiss LSM880 or Zeiss 980) using a 63x/1.4 objective lens. GFP was excited using a 488nm laser and emission collected from 495 to 500 nm and mCherry was excited using the 561nm laser and emission collected from 570 – 620 nm using an Airyscan (LSM880) or Airyscan 2 (LSM980) detector. Images were processed using Zeiss Zen software and modified (e.g. coloured) and quantified using ImageJ software (NIH).

### Recovery Growth Assays

Equivalent volumes of cells were harvested from mid-log cultures and washed three times with raffinose media before being resuspended in raffinose media and incubated in a shaking incubator at 30°C for 2 hours. Equivalent volumes of the raffinose starved cells were harvested and washed three times with glucose media before being resuspended in 100 μl of glucose media. This was added to 3 ml of glucose media and absorbance at OD_600_ was measured to obtain time point 0. Subsequent OD_600_ measurements were taken every hour using a plate reader (Thermo Scientific) and normalised to wild-type cells.

### Spot Growth Assays

Equivalent volumes of indicated cells from mid-log phase cultures were harvested and a 10-fold serial dilution was created in water prior to spotting on solid agar media plates. Plates were incubated at 30°C and images were captured at indicated time-points.

### Bioinformatic and Statistical analyses

Prediction of kinase consensus motifs in Pil1 was undertaken by submitting the Pil1 amino acid sequence to the yeast database in Netphorest (Horn et al., 2014) and experimentally determined phospho-sites (Luo et al., 2008; Walther et al., 2007) were selected for analysis. Raw data for analyses included in **Supplemental Table T1**. The results were filtered with a minimum phosphorylation probability score of 0.1. Unpaired Student’s *t*-tests were performed using GraphPad Prism v8.3.1. to compare the statistical significance between wild-type cells and mutants, or otherwise indicated pairwise comparisons, in described experimental conditions, with p values documented in **Supplemental Table T4**.

## ACKNOWLEDGMENTS

We would like to thank staff at the York Bioscience Technology Facility for technical assistance. We are very grateful to Robert Farese and Tobi Walther (Harvard Medical School) for providing us with antibodies raised against Pil1, to Andreas Mayer (University of Lausanne) for providing us with antibodies raised against Glc7, to Scott McIsaac (Calico Life Sciences, LLC) for sending the YETI yeast library used to modulate *GLC7* expression, and to Paul Pryor for access to the over-expression plasmid library used to over-express Pkh2. This research was supported by a Sir Henry Dale Research Fellowship from the Wellcome Trust and the Royal Society 204636/Z/16/Z (CM).

