## Supplementary material for "The phosphatase Glc7 controls eisosomal response to starvation via posttranslational modification of Pil1": Supp material

### SUPPLEMENTAL FIGURES AND LEGENDS

- **Figure S1:** Localisation phenotypes of Pkh kinases.
- **Figure S2:** Pil1 phosphorylation profiles of implicated kinase mutants.
- **Figure S3:** Pil1 is dephosphorylation in response to glucose starvation.
- **Figure S4:** Localisation of phosphatases at log and stationary phase.
- **Figure S5:** Optimising Glc7 expression and localisation tools.
- **Figure S6:** Quantification method for Pil1 localisation phenotypes.
- **Figure S7:** Growth assays of strains expressing Pil1-8A/8D and *glc7* mutants.
- **Figure S8:** Screening Glc7 regulatory subunits.

### SUPPLEMENTAL TABLES

- **Supplemental Table T1:** Netphorest hits
- **Supplemental Table T2:** Yeast Strains used in this study.
- **Supplemental Table T3:** Antibodies used in this study.
- **Supplemental Table T4:** Statistical analyses.

A

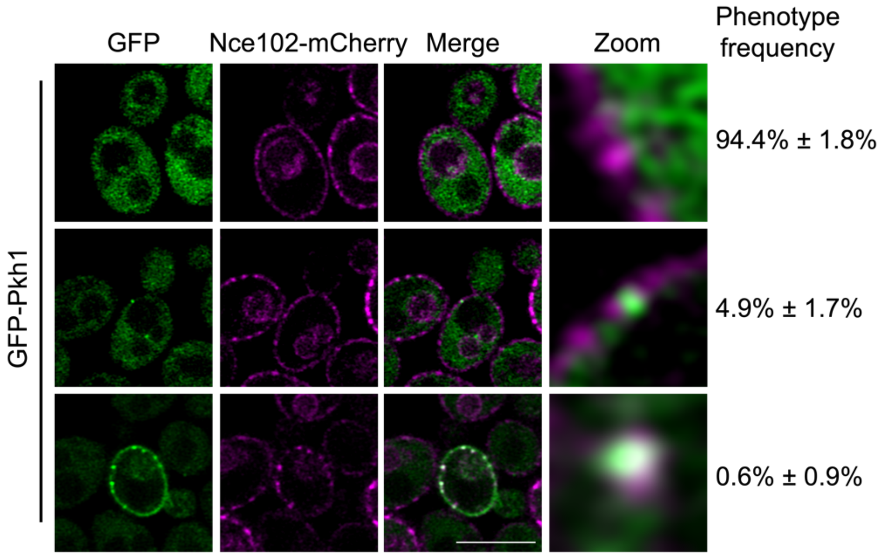


B

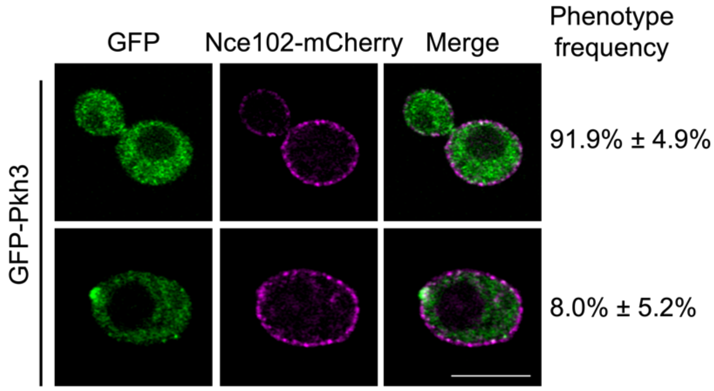


***Figure S1: Localisation phenotypes of Pkh kinases.***

**A - B)** Wild-type cells co-expressing Nce102-mCherry with either GFP-Pkh1 (**A**) or GFP-Pkh3 (**B**) were imaged using confocal Airyscan 2 microscopy and the frequency of each phenotype observed was quantified. Scale bar = 5μm.

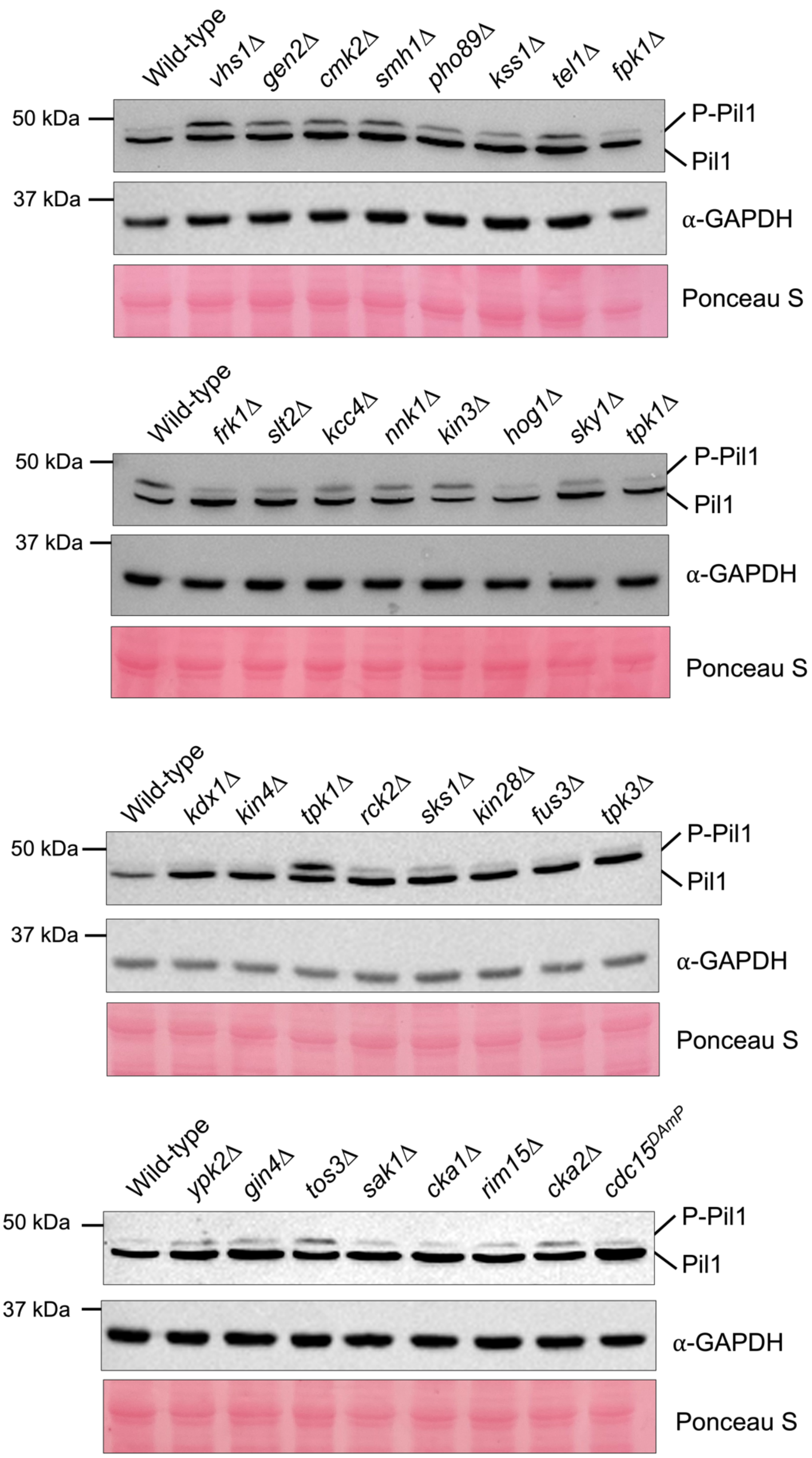


***Figure S2: Pil1 phosphorylation profiles of implicated kinase mutants.***

Wild-type and mutant cells either lacking non-essential kinases (Δ) or with reduced expression of essential kinases (*DAmP*) were grown to mid-log phase and lysates were generated for immunoblot using α-Pil1and α-GAPDH antibodies. Ponceau S stained membranes are also provided as loading controls.

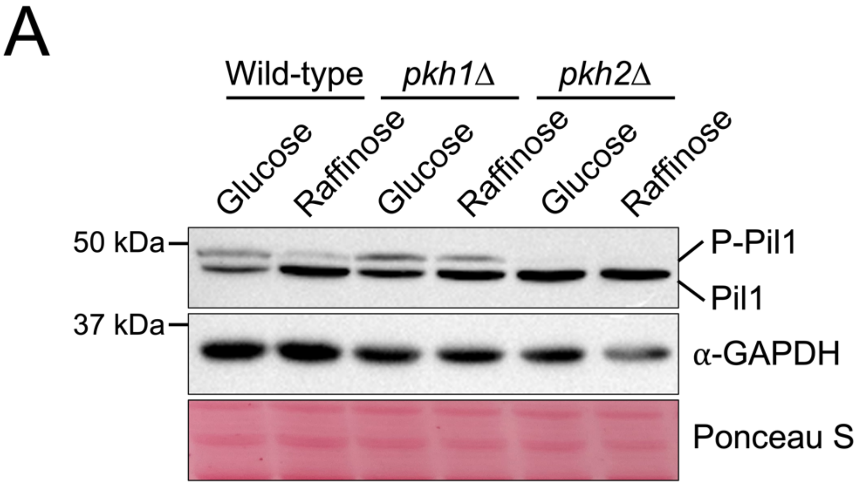


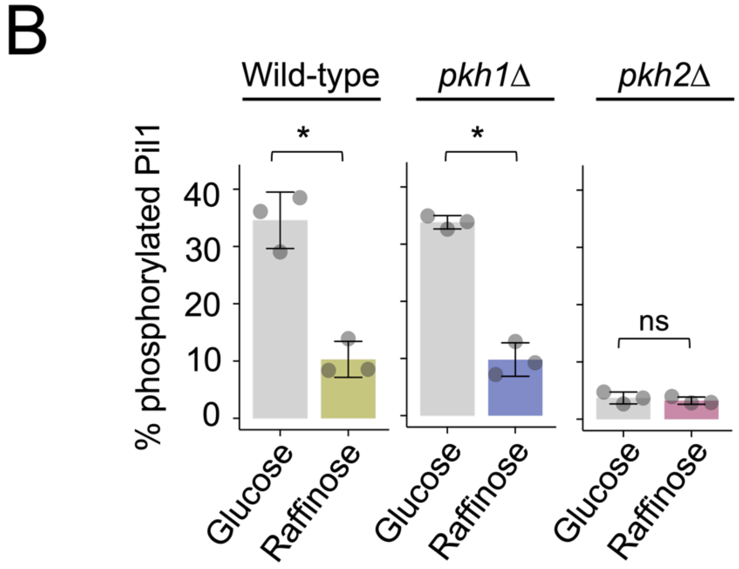


***Figure S3: Pil1 dephosphorylation in response to glucose starvation.***

**A)** Whole cell lysates of wild-type, *pkh1*Δ and *pkh2*Δ cells in glucose media and after 10 minutes of raffinose treatment were analysed by immunoblotting using α-Pil1 and α-GAPDH antibodies; Ponceau S stained membrane is shown. **B)** The percentage of phosphorylated Pil1 from each strain was quantified and statistical significance was determined using a Student's *t*-test.

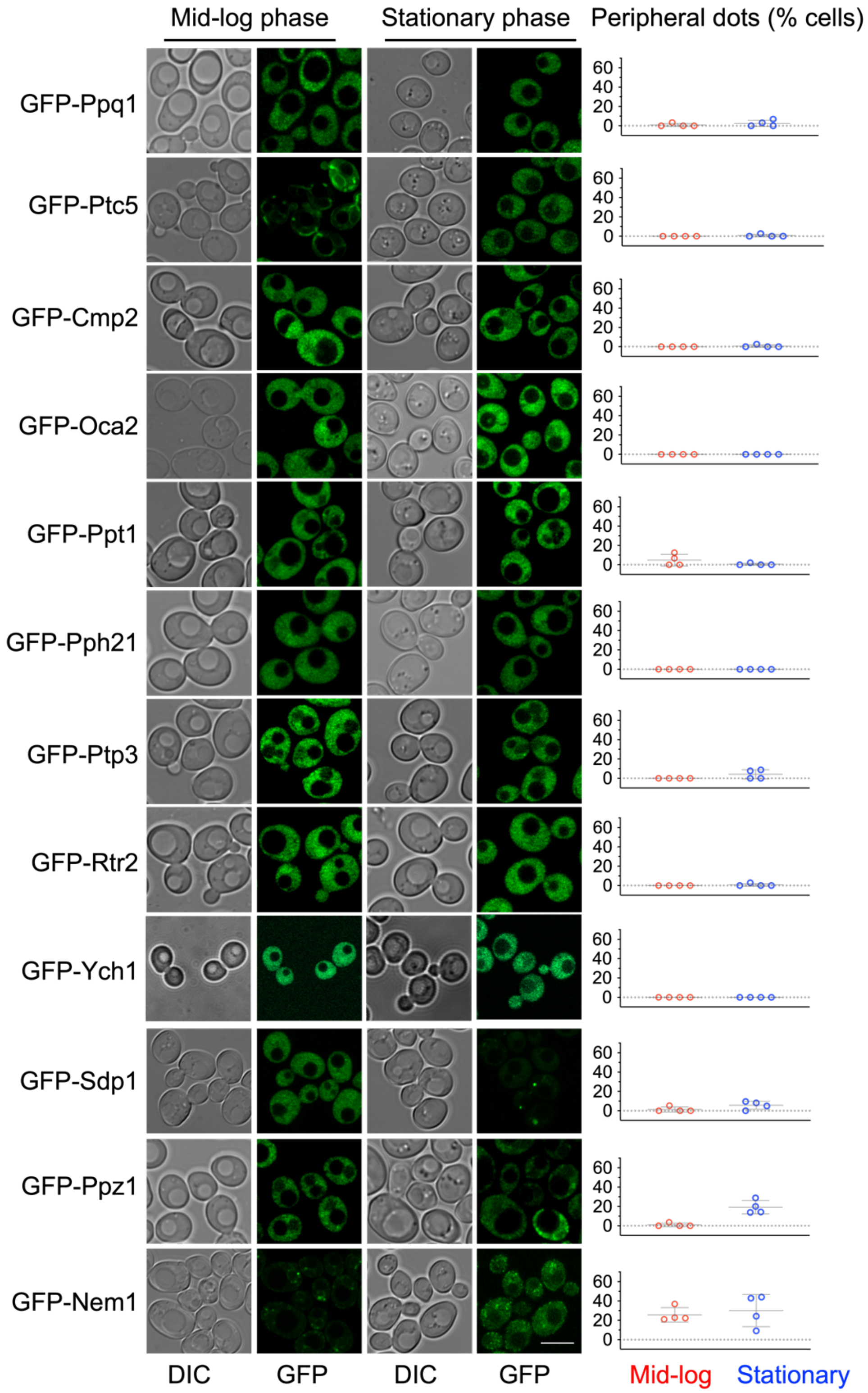


### *Figure S4: Localisation of phosphatases at mid-log and stationary phase.*

Indicated GFP tagged phosphatases were imaged using confocal Airyscan microscopy at mid-log and stationary phase. The number of peripheral dots per cell (n > 50) was quantified and the average for each experiment plotted (n = 4). Scale bar = 5µm.

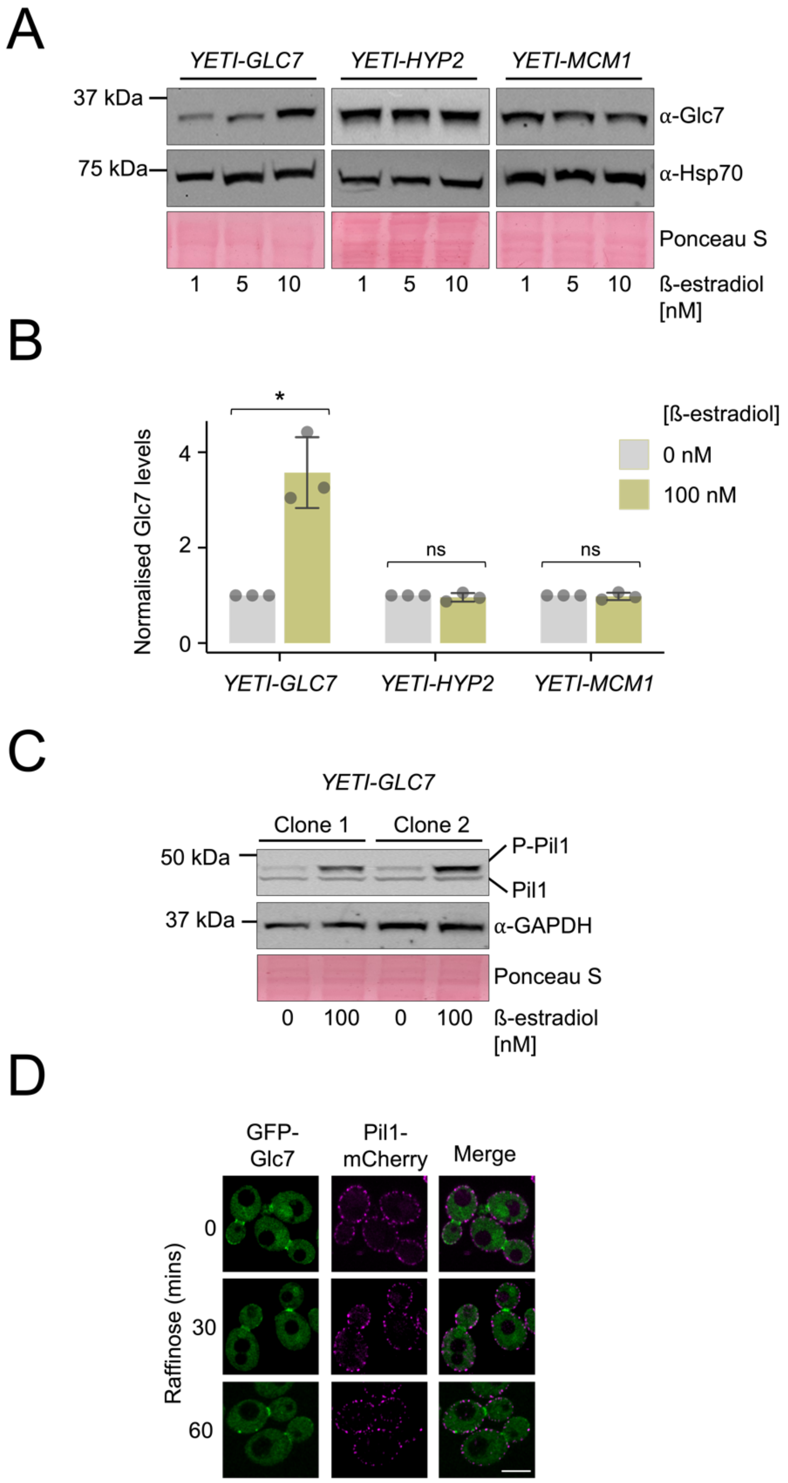


***Figure S5: Optimising Glc7 expression and localisation tools.***

**A)** Indicated strains were grown overnight in YPD media followed 6 hour incubation with 1 nM, 5 nM and 10 nM β-estradiol prior to the generation of whole-cell lysates and immunoblots using α-Glc7 and α-Hsp70 antibodies. **B)** Levels of Glc7 from immunoblots normalised to loading were quantified. (n=3) with and without β-estradiol. **C)** *YETI-GLC7* cells were grown overnight in 0 and 100 nM β-estradiol prior to whole-cell lysate generation and immunoblots using α-Glc7 and α-GAPDH antibodies. **D)** GFP-Glc7 and Pil1-mCherry were grown to mid-log phase and imaged using confocal microscopy at 0, 30 and 60 minutes of raffinose treatment. Scale bar = 5μm.

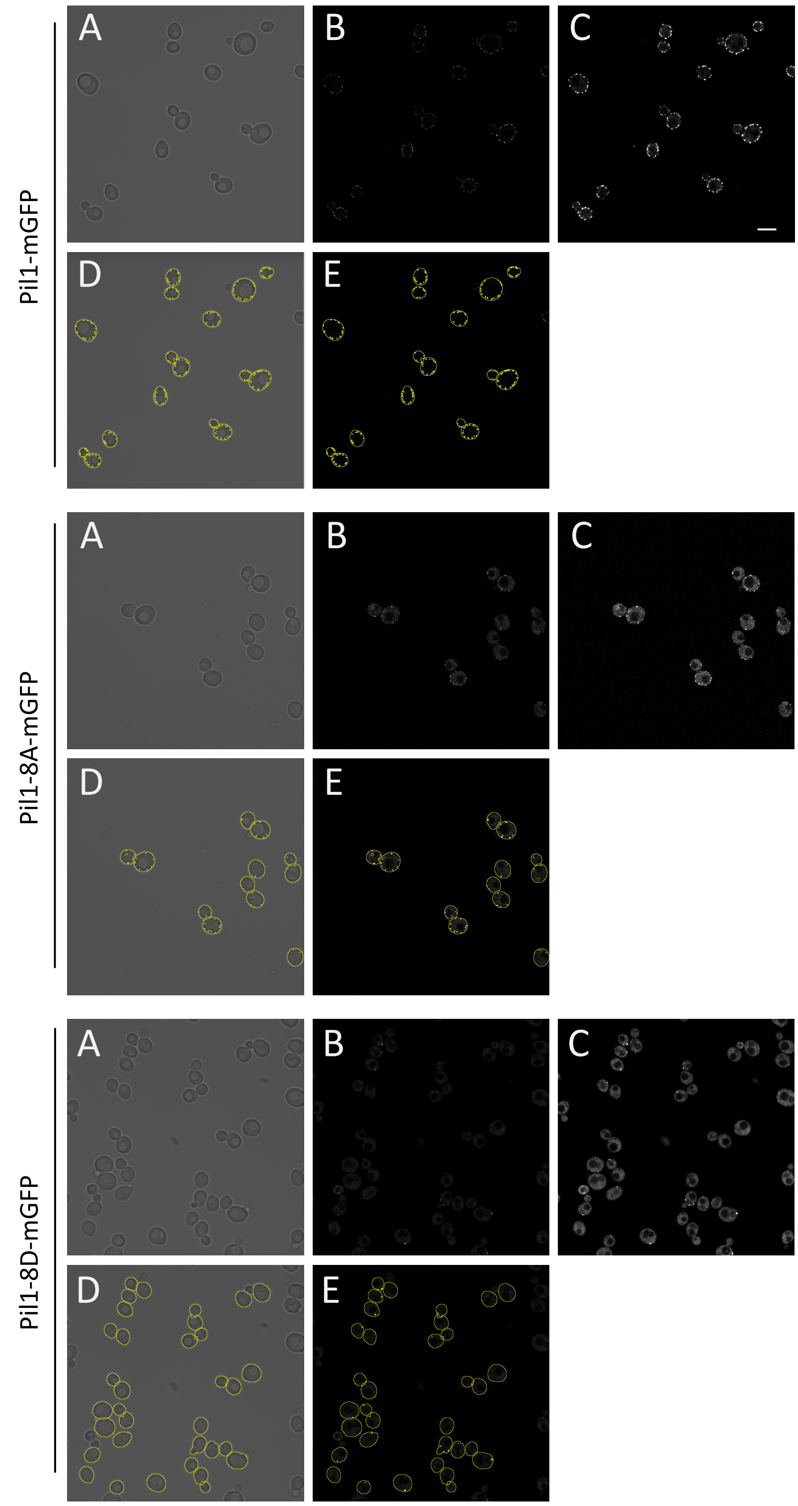


*Figure S6: Quantification method for Pil1 localisation phenotypes.*

Versions of Pil1 tagged with mGFP were imaged using Airyscan 2 microscopy, brightfield (A) and fluorescence channels (B) are shown. A brighter version of GFP signal is included (C) to better demonstrate localisation distribution between eisosomes and the cytoplasm but not used for quantifications. Whole cell segmentation (A) was performed using Cell Magic Wand tool and combined with eisosome-only segmentation (D) using *ostu* thresholding using ImageJ. The number of eisosomes per cell were counted and the percentage of eisosome signal compared to total cellular signal was calculated from these regions of interest from each cell. Scale bar = 5μm.

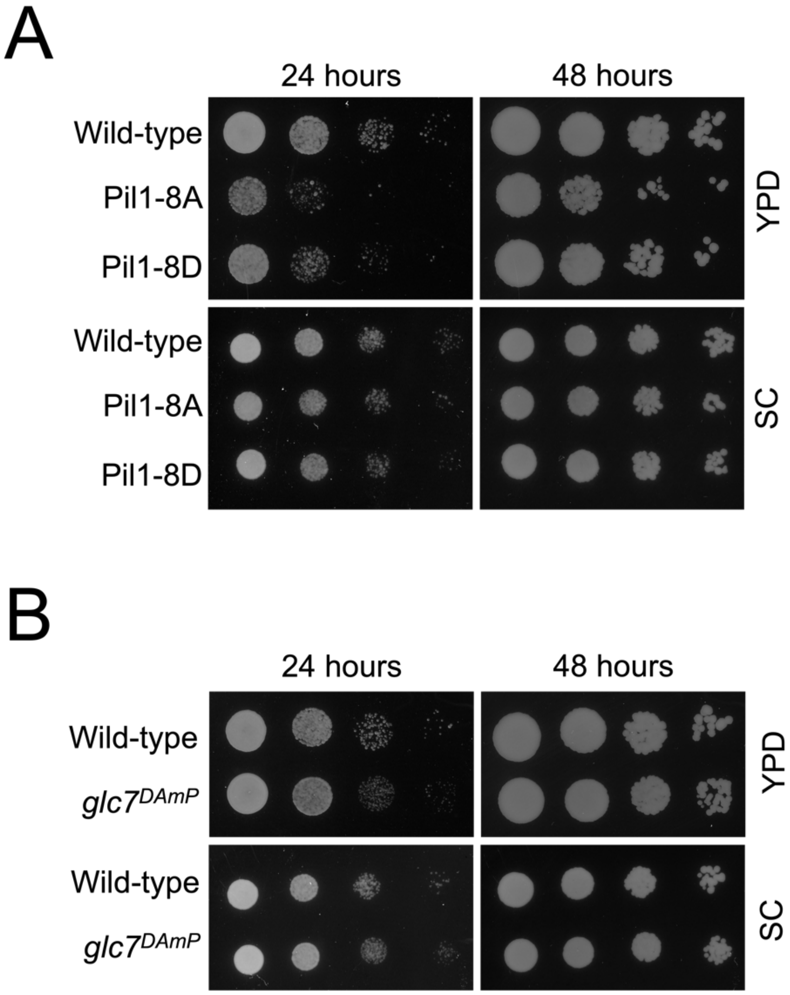


***Figure S7: Growth assays of strains expressing Pil1-8A/8D and glc7 mutants.***

**A – B)** Wild-type cellular growth was compared to either Pil1-8A and Pil1-8D expressing cells **(A)** or *glc7*$^{DAmP}$ mutants **(B)** by growing cells to mid-log phase in media (SC or YPD) before equivalent cell numbers were estimated by optical density measurements and harvested. 10-fold serial dilutions for each culture were generated, and yeast spotted out on both YPD plates and SC plates. Growth was recorded at 24 and 48 hours.

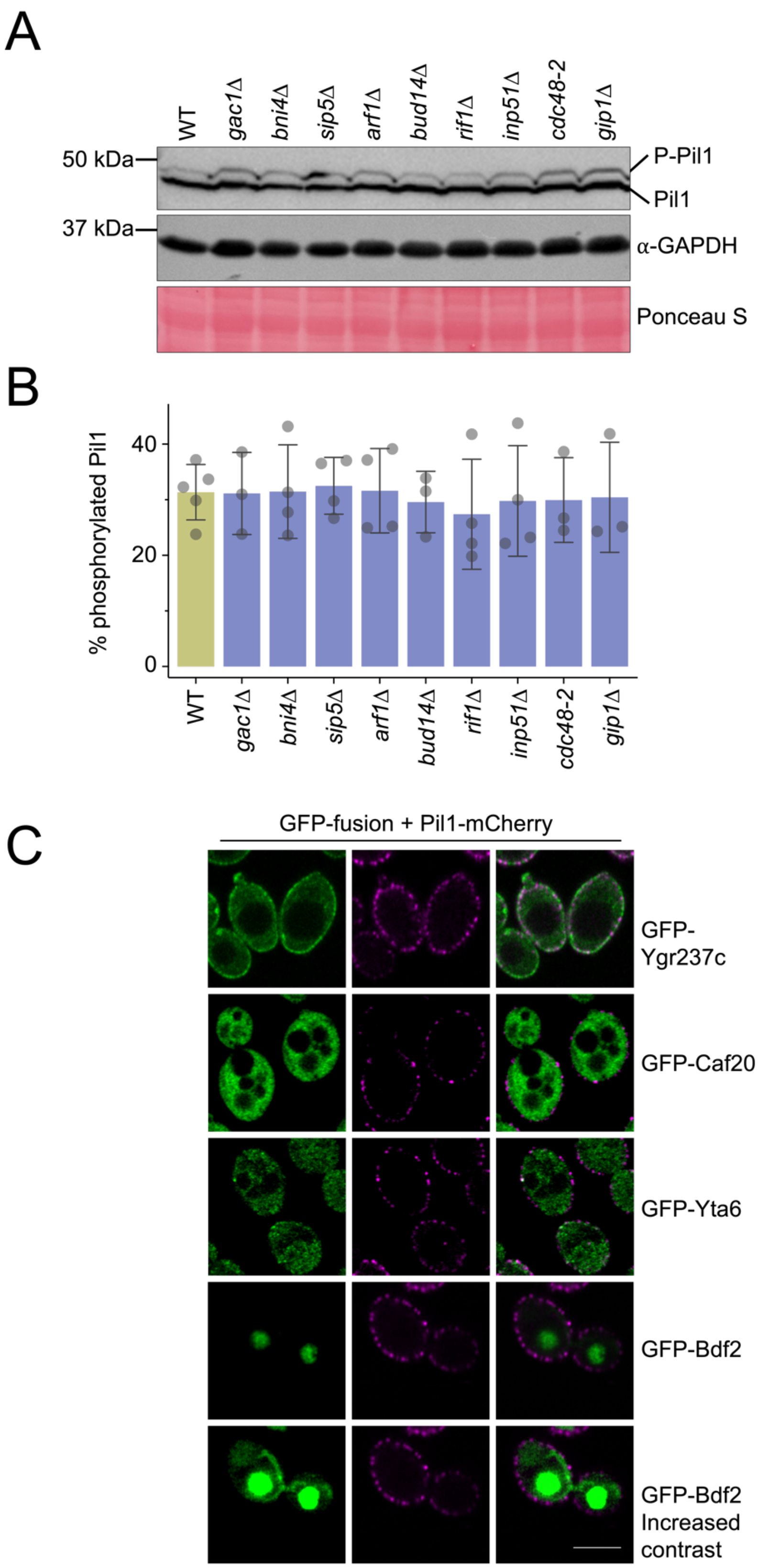


***Figure S8: Screening Glc7 regulatory subunits.***

**A)** Wild-type and mutant cells were grown to mid-log phase and lysates were generated for immunoblot using α-Pil1and α-GAPDH antibodies. Ponceau S stained membranes are also provided as loading controls. **B)** Immunoblot in **A** was repeated (n>3) and % of phosphorylated Pil1 species was quantified. Statistical significance was determined using a Student's *t*-test. **C)** Indicated GFP-fusion strains were co-expressed with Pil1-mCherry and imaged at mid-log phase using confocal microscopy (Airyscan 2) to determine localisation. Scale bar = 5μm.
